# Animal collocation analysis 2.0: improving statistical inference and applications for cohort comparisons

**DOI:** 10.64898/2026.03.20.713138

**Authors:** Elliot Howard-Spink, Monika Mircheva, Judith M. Burkart, Simon W. Townsend

## Abstract

Many animals communicate using sequences of signals, but identifying recurrent, non-random signal combinations remains methodologically challenging. Collocation analyses are increasingly popular approaches for detecting which signals animals combine at rates greater than expected by chance. However, existing methods for animal collocation analysis face several limitations that reduce their statistical rigour: they lack uncertainty estimates, fail to control for non-independence in sampled data, and do not account for inflated family-wise error rates when identifying attraction among many different signal types. These limitations restrict the broader applicability of animal collocation analysis, including preventing robust comparisons of signal combination strength between cohorts (e.g. populations, sexes or age classes). We adapt a novel form of Multiple Distinctive Collocation Analysis using Pearson residuals (MDCA-Pr) that addresses these statistical limitations, and validate its use in animal communication research in three ways: first, using numerous simulated datasets of different sizes and levels of signal recombination; second, using simulated data to evaluate the performance of MDCA-Pr in intercohort comparisons, and third, by demonstrating how MDCA-Pr can be applied to compare the vocal sequences produced by male and female captive-living common marmosets (*Callithrix jacchus*). MDCA-Pr shows high sensitivity, including at small sample sizes, and generally low false-positive rates, which we further reduce by applying additional criteria for identifying attraction between signals. During intercohort comparisons, MDCA-Pr is conservative, with low false-positive rates, and statistical power increases with sample size. MDCA-Pr is a robust method for evaluating signal attraction in animal communication and enables accurate intercohort comparison of animal signal combinations.

**Significance Statement:** By assessing the performance of MDCA-Pr on simulated animal-like data, we demonstrate that this method reliably detects signal combinations within and across animal cohorts, while overcoming statistical limitations of previous collocation analyses. We present an analytical pipeline for applying MDCA-Pr to animal signal data, including for intercohort comparisons, enabling identification and comparison of combinatorial strategies across entire signal repertoires. We illustrate this approach by comparing call combination strategies of male and female common marmosets when presented with food under experimental conditions, finding similar combinatorial strategies between sexes. MDCA-Pr therefore permits rigorous characterization of animal signal combinatoriality and opens avenues for investigating how demographic, social, and group-level factors influence combinatorial patterns.

## 1. Introduction

Growing research suggests that animals combine communicative signals into non-randomly structured sequences (herein, *signal combinatoriality*; Engesser and Townsend 2019; Suzuki and Zuberbühler 2019), with increasing examples being reported across species of birds (e.g. Engesser et al. 2016, 2024; Suzuki et al. 2016; Spiess et al. 2022; Walsh et al. 2024), and mammals (e.g. Collier et al. 2017, 2020; Gu et al. 2023; Hedwig and Kohlberg 2024), particularly primates (Schlenker et al. 2016a; Bosshard et al. 2024; Berthet et al. 2025; Girard-Buttoz et al. 2025; Le Floch et al. 2026). Signal combinatoriality has been proposed to enhance the flexibility of animal communication, as smaller repertoires of signal units (e.g. calls, gestures etc.) can be combined into a much broader repertoires of combinations (Nowak et al. 2000). In some instances, the sequential order of units within combinations is important for encoding information between signallers and receivers (Engesser et al. 2016, 2019; Suzuki et al. 2016; Engesser and Townsend 2019; Leroux et al. 2023; Berthet et al. 2025; Girard-Buttoz et al. 2025); for example, wild chimpanzees (*Pan troglodytes*) have been documented producing ‘Grunt→Hoo’ combinations frequently in travelling and fusion contexts, whereas the reverse ‘Hoot→Grunts’ are used when stopping to feed and rest (Girard-Buttoz et al. 2025). Characterizing how animals combine signals is therefore crucial for understanding the structural flexibility and potential function of animal communication (Schlenker et al. 2016b&c, 2024). In addition, comparative research into animal signal combinatoriality represents one of a handful of avenues for understanding the evolution of human languages’ extensive combinatoriality, including humans’ dual generative systems of phonology and lexical syntax (Collier et al. 2014; Coye et al. 2018b; Fitch 2018; Townsend et al. 2018; Idsardi 2019; Zuberbühler 2019; Leroux and Townsend 2020).

Evaluating signal combinatoriality in animal communication systems requires robust methods to quantify the relative frequencies with which signals are combined, and whether these frequencies exceed those expected by random co-occurrence of signals over time. Derived from computational linguistics, collocation analysis represents one such method, and identifies whether specific pairs of sequence units (bigrams) co-occur more frequently (indicating ‘attraction’) or a less frequently (‘repulsion’) than expected based on the independent distribution of either signal type within a corpus of sample data. While several different forms of collocation analyses have been developed for analysis of human language corpora (Church et al. 1991; Stefanowitsch and Gries 2003, 2005; Bartsch 2004; Nesselhauf 2005; Xiao and McEnery 2006; Evert 2009; Lehecka 2015), Bosshard et al. (2022) validated two forms of collocation analysis for analysing signal combinations within animal datasets (henceforth, ‘animal collocation analysis’; methods include Multiple Distinctive Collocation Analysis: MDCA, & Mutual Information Collocation Analysis: MICA). These forms of animal collocation analysis allowed researchers to transition from qualitative assessment to quantitative investigation of combinatoriality in animal signal repertoires. Consequently, several studies have applied animal collocation analysis to identify significantly attracted signal combinations within animal vocalizations (Leroux et al. 2021, 2022), multimodal signals (Wewhare and Krishnan 2024; Mine et al. 2024, 2025), and during dyadic interactions (Demartsev et al. 2024; van Boekholt et al. 2025), as well as in the pairwise structure of non-communicative sequential behaviours, such as animals’ food item pairings during foraging (Freymann et al. 2024). However, while animal collocation analysis has proven to be a valuable tool for identifying re-occurring pairwise patterns in animals’ sequential behaviours, current forms of animal collocation analysis face several limitations that potentially compromise the validity of their outcomes (see Gries 2022):

Firstly, existing methods for animal collocation analysis only yield point estimates of collocation strength, and do not provide measures of uncertainty surrounding collocation strength estimates, such as confidence intervals (limitation 1). In the absence of these measures, it is not possible to evaluate the precision of estimated collocation strength, and this prevents thorough examination of the strength of evidence for attraction or repulsion of signal combinations.

Secondly, all animal studies to date have reported significant attraction between signals using minimal collocation-strength thresholds (‘pbins’, based on log-transformed binomial probability scores or mutual information; Bosshard et al. 2022). However, no methods have been suggested for how to control for inflated family-wise error rates when comparing increasingly more signal types within these analyses (increasing the likelihood of type 1 errors for significant attraction or repulsion; limitation 2).

Thirdly, while existing methods for collocation analysis compare the frequency of signal co-occurrence relative to what is expected if each signal type was distributed independently in a sample, they do not account for whether nested relationships exist within sample data which could also influence the frequency of signal co-occurrence (limitation 3). For example, pairwise signal transitions may be recorded from the same bout, or same individual, meaning signal bigrams sampled for the analysis are not independent. Currently, no methods have been proposed regarding how to account for this non-independence when generating point-estimates in animal collocation analysis.

Finally, current methods for animal collocation analysis impose a trade-off between identifying directionality in collocates (using MCDA, where A→B differs to B→A) or to maintaining higher sensitivity to rarer but significant collocates (MICA, which is blind to the direction of transitions; Bosshard et al. 2022; limitation 4). Analyses that can combine sensitivity and directionality are therefore needed, as directionality can be a functionally-meaningful aspect of animals’ signal combinations (Girard-Buttoz et al. 2025).

In an effort to overcome these obstacles when analysing human language corpora, Gries (2023) recently proposed several adaptations to MDCA to improve the analysis’ statistical rigour. Classical MDCA involves iterative construction of 2×2 binomial arrays to estimate the probability of each type of sequence unit being combined, and thus its collocation strength; however, this is computationally costly, and can be time consuming. Therefore, Gries proposes quantifying collocation strength using Pearson residuals from a singular array tallying all pairwise unit co-occurrences. Pearson residuals (*Pr*) are scaled comparisons of the number of instances a particular pair of units co-occurred (*O*) compared to the number of times is Expected (*E*) by chance given the frequency of either sequence unit type in the sample corpus (Liao et al. 2024):

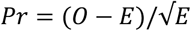

By estimating these residuals from one global table, MDCA by Pearson residuals (MDCA-Pr) considerably reduces the computational burden of collocation analysis. This lends the analysis to bootstrapping, where uncertainty estimates for the collocate strength for each pair of signal units can be generated, e.g. confidence intervals for collocation strength (solving limitation 1; see Gries 2023). The size of confidence intervals can be adjusted through Bonferroni correction, thus accounting for family-wise error rates that have been inflated by multiple comparisons of different signal pairs (solving limitation 2; see Studies 1 & 2 below). Moreover, through block bootstrapping (resampling with replacement at highest hierarchical levels in the corpus data, e.g. by resampling all bouts produced by a particular individual all at once), uncertainty measures for collocation strength can be estimated whilst accounting for nested structure in the data (solving limitation 3; see Gries 2023). As MDCA-Pr is also directional, this method may have the potential to balance directionality and sensitivity (solving limitation 4). However, the performance of MDCA-Pr has yet to be tested on the types of datasets collected by animal behaviour researchers, which are often small, skewed, and contain far fewer element types than linguistic corpora.

In addition to these statistical improvements, MDCA-Pr may offer new avenues to explore questions that are intractable with previously validated forms of animal collocation analysis, including whether signal combinatoriality can be compared between cohorts of animals (e.g. different groups or populations of animals, or different sex or age classes within a species). Emerging studies indicate that for some species, signal combinations may differ between cohorts (usually animal populations; Schlenker et al. 2014; Girard-Buttoz et al. 2022a), as may the contexts in which shared signal combinations are used (Schamberg et al. 2024), possibly reflecting cohort-specific communicative norms. However, these studies have overwhelmingly focused on the *rates* with which signals are combined (irrespective of the signal units) or the presence/absence of specific signal combinations. As of yet, no studies have characterized the extent of intraspecific variation in combinatoriality across a species’ entire signal repertoire, largely because of the limited availability of analytical tools capable of supporting such comparisons. Previously validated approaches to collocation analysis cannot fill this methodological gap, as they do not provide uncertainty estimates for collocation strength. This makes it impossible to assess whether observed differences between cohorts reflect valid behavioural differences, or if they are the product of trivial sampling error. However, because MDCA-Pr can be easily bootstrapped to estimate confidence intervals, it may permit formal testing for significant differences in collocation strength between cohorts. Given this potential, the use of MDCA-Pr during intercohort comparison would also benefit from empirical validation.

Here, we take a three-pronged approach to validate the suitability of MDCA-Pr to analysing animal behavioural data, including during intercohort comparison, using a mixture of synthetic vocal data and vocal data taken from captive-living common marmosets (*Callithrix jacchus)*:

In Study 1, we evaluate the performance of MDCA-Pr on synthetic animal-like vocal datasets of different sizes and levels of recombinatoriality, following a validation procedure from Bosshard et al. (2022; see Section 2.1). We quantify the sensitivity (true-positive rate) and selectivity (false-positive rate) of MDCA-Pr to evaluate its performance when identifying call attraction across these different types of animal datasets.

In Study 2, we evaluate the performance of MDCA-Pr when comparing synthetic vocal data from simulated animal cohorts (Section 2.2). We assess the sensitivity with which MDCA-Pr can identify pre-programmed differences in the probability of calls being combined between cohorts (the true-positive rate), and selectivity of the analysis, including whether any call combinations with the same probabilities between cohorts were erroneously identified as having different collocation strengths (the false-positive rate). Similarly to Study 1, we evaluate MDCA-Pr during intercohort comparison on synthetic datasets of varying sizes and levels of recombinatoriality, to build a comprehensive picture of the performance of MDCA-Pr given the different types of datasets that animal behaviour researchers frequently encounter.

Finally, in Study 3, we extend beyond simulated data, by applying MDCA-Pr to compare combinatorial strategies of captive-living male and female common marmosets in a feeding context (Section 2.3). This study allows us to illustrate how MDCA-Pr can be applied to data from real interacting animals in a controlled foraging context.

Together, this suite of analyses allows us to compare the performance of MDCA-Pr to previously-validated forms of animal collocation analysis, and to evaluate its efficacy in identifying similarities and differences in combinatorial behaviours between cohorts of animals. Additionally applying MDCA-Pr to marmoset call data demonstrates its applicability to real-world data, and provides an initial showcase of how cohort-level combinatoriality can be compared.

## 2. Methods & Results

Across all three studies, MDCA-Pr was performed using the method proposed by Gries (2023), using an adapted code in R (v. 4.5.1; R Core Team 2022; for original code see Gries 2024). In each study, the simulated and real call sequences were parsed into bigrams that described pairwise co-occurrence of calls. Each unique type of bigram was tallied in a global table, and Pearson residuals were calculated for each bigram type using the chisq.test() function from the stats package (base R v 4.5.1; R Core Team 2022). Pearson residuals were calculated across 10,000 bootstraps of the data (blocked at the level of the individual), and confidence intervals were estimated from this bootstrapped distribution for each bigram type, where the original 95% confidence interval (alpha = 0.05) was adjusted via Bonferroni correction to the number of comparisons being made (alpha_adjusted_ = 0.05/Number of bigram types compared; comparisons were counted as either each instance a bigrams’ collocation strength was compared to zero for a particular dataset (e.g. Study 1), or each instance a bigrams’ collocation strength was compared between two cohorts (e.g. Study 2)). Confidence intervals were estimated non-parametrically using upper and lower quantiles of the bootstrapped distribution, and were constructed for two-tailed comparison (lower percentile = 100 x alpha_adjusted_/2; upper percentile = 100 x (1 – alpha_adjusted_/2)). All scripts to run MDCA-Pr on animal data can be found in our supplementary code.

### 2.1 Study 1 – validation of MDCA-Pr using simulated data

#### 2.1.1 Simulating animal-like call data

Following Bosshard et al. 2022, we simulated four distinct datasets of animal-like calls. Two *Small* datasets (N_Individuals_ = 10; 16 bigrams per individual; total bigrams = 160), and two *Large* datasets (N_Individuals_ = 20; 160 bigrams per individual; total bigrams = 3200). For simplicity, we assume that each bigram comes from its own bout of 2 calls (i.e. for the *Small* dataset, 32 individual calls were produced in 16 pairs per individual) and provide an example of how to analyse longer call bouts in Study 3 (Section 2.3).

For each size category, an *Exclusive* dataset was simulated that only contained four call types combined into three distinct focal call bigrams: Peep→Howl, Howl→Peep and Huff→Puff (see Table 1). The *Small* and *Large Exclusive* datasets functioned as a positive control, where calls within focal bigrams should be identified as ‘attracted’ as no other competing call bigrams appear in the data.

**Table 1.** Simulated data and results of Study 1. All probabilities are rounded to 3.d.p, and collocation strengths presented to 3.s.f. Focal call bigram types are in bold. Bigram probability indicates the likelihood that each type of bigram is simulated during the production of data for each artificial individual. Mean collocate strengths are provided for focal call bigram types, alongside confidence intervals. Collocate strengths for all other bigram types are reported in Tables S1-S4. Asterisks indicate significant attraction for a given bigram type, including after applying the 1SD filter. Horizontal dashes indicate no significant attraction. For control datasets, no attraction was detected even prior to applying the 1SD filter

| Call Bigram type | Bigram Probability | MDCA-Pr Result |  |  |  |  |  |  |  |
| --- | --- | --- | --- | --- | --- | --- | --- | --- | --- |
|  |  | Small Dataset |  |  | Large Dataset |  |  | Control Datasets |  |
|  |  | Mean Collocate Strength | 1SD Upper Threshold | Attraction? | Mean Collocate Strength | 1SD Upper Threshold | Attraction? | Small | Large |
| **Peep→Howl** | **0.313** | 8.61 | 6.32 | * | 39.2 | 28.3 | * | - | - |
| **Howl→Peep** | **0.438** | 7.12 | 6.32 | * | 31.4 | 28.3 | * | - | - |
| **Huff→Puff** | **0.250** | 9.57 | 6.32 | * | 42.5 | 28.3 | * | - | - |
| Recombinatorial |  |  |  |  |  |  |  |  |  |
| **Peep→Howl** | **0.102** | 4.81 | 1.55 | * | 16.9 | 5.99 | * | - | - |
| *Peep→(Not Howl)* | 0.020 |  |  | - |  |  | - | - | - |
| **Howl→Peep** | **0.143** | 3.84 | 1.55 | * | 18.5 | 5.99 | * | - | - |
| *Howl→X (Not Peep)* | 0.020 |  |  | - |  |  | - | - | - |
| **Huff→Puff** | **0.082** | 1.94 | 1.55 | - | 14.8 | 5.99 | * | - | - |
| *Huff→X (Not Puff)* | 0.020 |  |  | - |  |  | - | - | - |
| *X→X (Any other combination, e.g. Hum→Cough)* | 0.020 |  |  | - |  |  | - | - | - |

Additionally, a *Recombinatorial* dataset was produced for each size category. These *Recombinatorial* datasets included the same three focal call bigram types as before (each with a different probability of occurring in the data; see Table 1). However, the *Recombinatorial* datasets also featured three additional call types (Cough, Whistle and Hum), which, alongside Peep, Howl and Puff, were programmed to flexibly combine into bigram types at lower probabilities (2% each). Some of these background combinations included call types present in the focal call bigrams (e.g. Peep→Cough, or Hum→Puff), whereas others did not (e.g. Cough→Hum). As the focal call bigrams occur at higher frequencies than other bigram types, focal call bigrams should be identified as attracted call combinations (as found by other forms of animal collocation analysis when run on data with the same bigram probability distributions; Bosshard et al. 2022). In general, across the simulated datasets (*Small Exclusive*, *Small Recombinatorial*, *Large Exclusive*, *Large Recombinatorial*), those that are smaller and more recombinatorial present a greater challenge for MDCA-Pr to identify attraction between calls in the focal bigrams.

In addition, we ran MDCA-Pr on four datasets that were of equivalent size to each of these four datasets and had the same numbers of call types; however, call bigrams were constructed by sampling call pairs from a uniform random distribution. These randomized datasets offered a negative control, where no significant attraction or repulsion should be identified between any call types.

#### 2.1.2 Evaluating MDCA-Pr performance

MDCA-Pr was applied to each dataset independently. Following existing literature, we only refer bigrams as call combinations if they are identified as experiencing significant levels of attraction (e.g., Leroux et al. 2022). To identify attracted call combinations across all bigram types, we followed a two-step procedure:

1. To identify call combinations that was unlikely to be driven by chance association in the data, we initially identified attraction by determining which call bigram types had confidence intervals for collocate strength that were wholly above zero.
2. After evaluating the results from step 1, we extended this initial assessment of attraction by following a recent study that applied MDCA to animal communication data (van Boekholt et al. 2025). This study reported that MDCA can identify many low-level attracted combinations in behavioural units, potentially to frequencies that suggest instances false-positive attraction. The authors proposed using an additional threshold to identify ‘strongly attracted’ bigrams, where attracted call combinations required a collocation strength that surpassed two standard deviations of collocation strengths for all bigram types. We evaluated whether a similar threshold would enhance the performance of MDCA-Pr using a post-hoc analysis, where we reclassified bigram types as ‘attracted’ if their mean collocate strength was one standard deviation higher than the mean of all collocate strengths. We chose to start at one standard deviation (rather than two) to test the methodological benefits of a less conservative threshold for MDCA-Pr when accounting for false-positive rates. If false-positive rates persisted under this threshold, our analysis lends itself to calibration of a more suitable threshold that can address low-level false-positive rates across datasets of different sizes and levels of recombinatoriality.

For each simulated dataset, we evaluated the performance of MDCA-Pr using measures of sensitivity and selectivity. Sensitivity was measured by the number of true positives identified (more means higher sensitivity), with a true positive being any instance where significant attraction was identified for the focal call bigram types. Selectivity was measured by the number of false positives identified (lower means better selectivity), with false positives categorized as any instance where significant attraction was detected between non-focal call bigram types. We focused on attraction only, and did not examine significant repulsion between call types (Gries 2023), as animal communication studies almost exclusively aim to characterize which calls animals frequently combine, rather than those animals rarely or never combine.

#### 2.1.3 Results

When identifying attracted call combinations solely based on the confidence intervals for collate strength, MDCA-Pr demonstrated perfect selectivity and sensitivity across both *Small* and *Large Exclusive* datasets. All three focal call bigram types were identified as significantly attracted, and no false-positive attractions were identified (see Fig. 1. & Table 1; see Tables S1 & S2). As predicted, non-simulated combinations (e.g. Peep→Puff) were significantly repulsed as these call pairings never occurred within the data.

**Figure 1.**
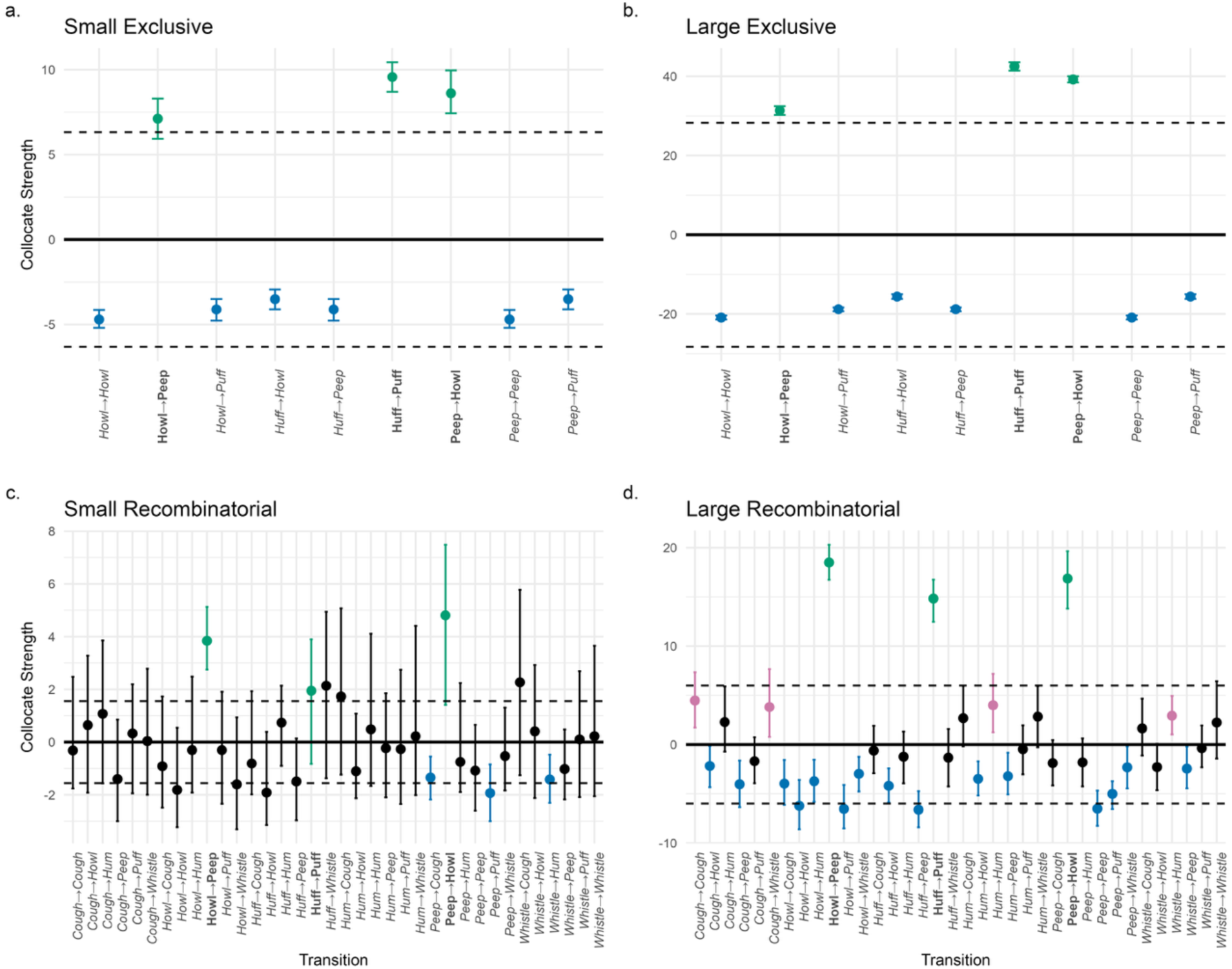
MDCA-Pr on the four non-random simulated datasets from Study 1. Points indicate the mean collocation strength for a given call bigram type across bootstraps, and error bars indicate the confidence interval surrounding each mean (95% CI adjusted by Bonferroni correction; see Section 2). Focal call bigrams labelled in bold and plotted in green. Data plotted in blue indicates repulsion between two call types (CI wholly below zero). Data plotted in red indicates initial attraction for non-focal call bigrams (CI wholly above zero) prior to applying the +-1SD filter (indicated by the dashed horizontal lines). Zero is indicated by a solid horizontal line

Perfect selectivity was achieved for the *Small Recombinatorial* dataset, as no false-positive attraction was identified. However, sensitivity was somewhat lower for the *Small Recombinatorial* dataset: only two focal call bigram types were identified as significantly attracted (Peep→Howl, & Howl→Peep; Fig. 1; Tables 1 & S3). For Huff→Puff, the focal call bigram type with the lowest combination probability, the confidence interval for collocation strength overlapped zero.

Increasing the size of the *Recombinatorial* data improved sensitivity, as in the *Large Recombinatorial* dataset, all focal call bigram types were identified as attracted (see Fig. 1 & Table S4). However, this came at the expense of selectivity, as significant lower-level attraction was identified between four of the non-focal call bigram types.

For each of the four datasets, we reclassified ‘attraction’ as any instance where a call bigram type had a confidence interval for collocation strength that was wholly above zero, and a mean collocation strength that exceeded one standard deviation of the collocation strengths across all bigram types. Following this reclassification, the sensitivity of our analysis was unchanged, with attraction being identified at the same rate as before across all focal bigram types. However, selectivity was improved after this threshold was applied, with no false-positive attraction being identified in any of the datasets (see Fig. 1 & Table 1).

In the *randomized control* datasets, no significant attraction or repulsion was detected (both with and without the addition of a 1SD filter), meaning MDCA-Pr is unlikely to identify significantly attracted call combinations in completely randomized data (see Table 1 & Fig. S1).

### 2.2 Study 2 – intercohort comparisons using simulated data

#### 2.2.1 Simulating call bigram data across cohorts

To evaluate whether MDCA-Pr could be used to compare combinatoriality across cohorts of animals, we simulated four classes of dataset, each containing three corpora of call bigrams. These corpora were of a fixed size within each class (*Small* & *Large*; following the size categorizations from Study 1) and contained specific levels of recombinatoriality (see below). For each class, two corpora were generated using the same probability distribution for all call bigram types (*Equivalent* cohorts 1 & 2) and a single corpus generated from a different probability distribution (an *Unequivalent* cohort).

Two classes of datasets were *Recombinatorial*: one *Large* and one *Small*. For the *Equivalent Recombinatorial* corpora in each class, we used the same probabilities for each call bigram type as the *Recombinatorial* datasets used in Study 1. Whereas, for the *Unequivalent Recombinatorial* corpus in each class, we reduced the probability of each focal bigram type occurring in the data by increasing amounts (Peep→Howl: −5%; Huff→Puff: −7.5%; Howl→Peep: −10%; see Table 2). For each reduction in the probability of a focal call bigram occurring in the data, all alternative transitions away from the first call in the bigram received a small uniform increase in their likelihood of being produced (e.g., as Peep→Howl was reduced by −5%, the bigrams Peep→Cough, Peep→Hum, Peep→Peep, Peep→Puff, & Peep→Whistle all increased by 1% each). This ensured that the overall probability distribution across bigram types always summed to 1. Whilst many bigram types therefore have different probabilities of occurring in the *Equivalent* and *Unequivalent* corpora, we retain our use of ‘focal bigram type’ for Peep→Howl, Howl→Peep, and Huff→Puff, as these bigram types exhibit the greatest differences between the two types of corpora.

**Table 2.**
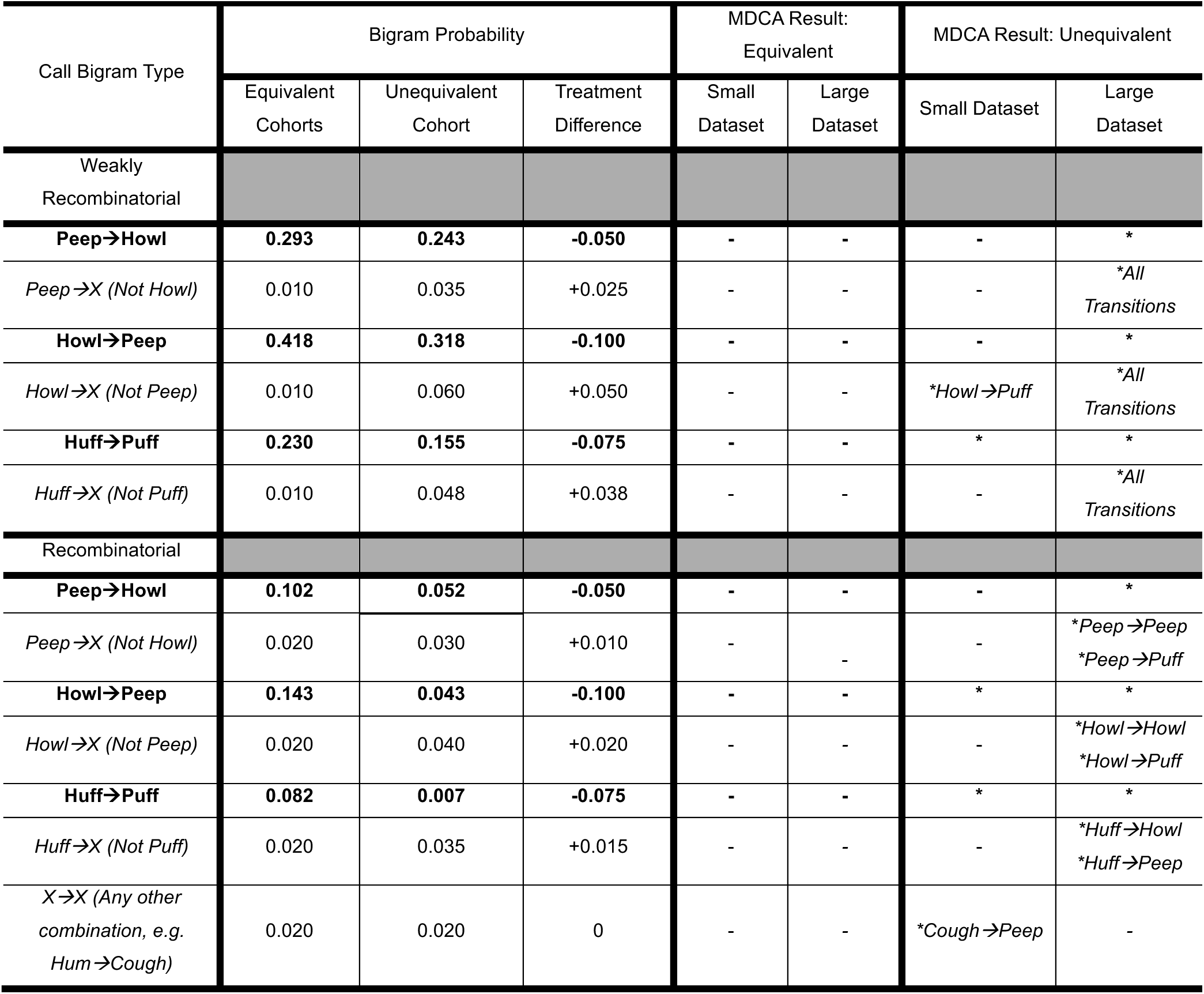
Simulated data and results of Study 2. Bigram probability indicates the likelihood that each type of bigram is simulated during the production of each dataset. Asterisks indicate that a significant difference in collocation strength between cohorts for a particular bigram type. Horizontal dashes indicate no significant difference. Specific values for collocate strengths for all call bigram types are reported in Tables S5-S8

| Call Bigram Type | Bigram Probability |  |  | MDCA Result: Equivalent |  | MDCA Result: Unequivalent |  |
| --- | --- | --- | --- | --- | --- | --- | --- |
|  | Equivalent Cohorts | Unequivalent Cohort | Treatment Difference | Small Dataset | Large Dataset | Small Dataset | Large Dataset |
| Weakly Recombinatorial |  |  |  |  |  |  |  |
| **Peep→Howl** | **0.293** | **0.243** | **-0.050** | - | - | - | * |
| *Peep→X (Not Howl)* | 0.010 | 0.035 | +0.025 | - | - | - | *All Transitions |
| **Howl→Peep** | **0.418** | **0.318** | **-0.100** | - | - | - | * |
| *Howl→X (Not Peep)* | 0.010 | 0.060 | +0.050 | - | - | *Howl→Puff | *All Transitions |
| **Huff→Puff** | **0.230** | **0.155** | **-0.075** | - | - | * | * |
| *Huff→X (Not Puff)* | 0.010 | 0.048 | +0.038 | - | - | - | *All Transitions |
| Recombinatorial |  |  |  |  |  |  |  |
| **Peep→Howl** | **0.102** | **0.052** | **-0.050** | - | - | - | * |
| *Peep→X (Not Howl)* | 0.020 | 0.030 | +0.010 | - | - | - | *Peep→Peep<br>*Peep→Puff |
| **Howl→Peep** | **0.143** | **0.043** | **-0.100** | - | - | * | * |
| *Howl→X (Not Peep)* | 0.020 | 0.040 | +0.020 | - | - | - | *Howl→Howl<br>*Howl→Puff |
| **Huff→Puff** | **0.082** | **0.007** | **-0.075** | - | - | * | * |
| *Huff→X (Not Puff)* | 0.020 | 0.035 | +0.015 | - | - | - | *Huff→Howl<br>*Huff→Peep |
| *X→X (Any other combination, e.g. Hum→Cough)* | 0.020 | 0.020 | 0 | - | - | *Cough→Peep | - |

We did not simulate any *Exclusive* classes of datasets, as any reductions in the probability of a focal call bigram occurring in the data had to be compensated for by increasing the probability a related call bigram being sampled during data simulation (i.e. a reduction in A→B led to an increase in A→C), thus we were not able to maintain the exclusivity property when simulating an *Unequivalent* corpus.

Instead, we simulated two *Weakly Recombinatorial* classes of datasets, which had the same call types as those in the *Exclusive* datasets from Study 1 (Howl, Peep, Huff & Puff), but in addition to the three types of focal call bigrams, these calls were recombined into several non-focal bigram types at low levels (each with a 1% probability of occurring; see Table 2 & Fig 2). As before, we simulated *Small Weakly Recombinatorial* and *Large Weakly Recombinatorial* classes of data, each with two *Equivalent* corpora, and one *Unequivalent* corpus. For the *Unequivalent* corpus, the probability of each focal call bigram occurring in the data was reduced by the same amounts as in the *Recombinatorial* classes, and these probabilities were distributed among other bigram types in the same way (see Table 2).

**Figure 2.**
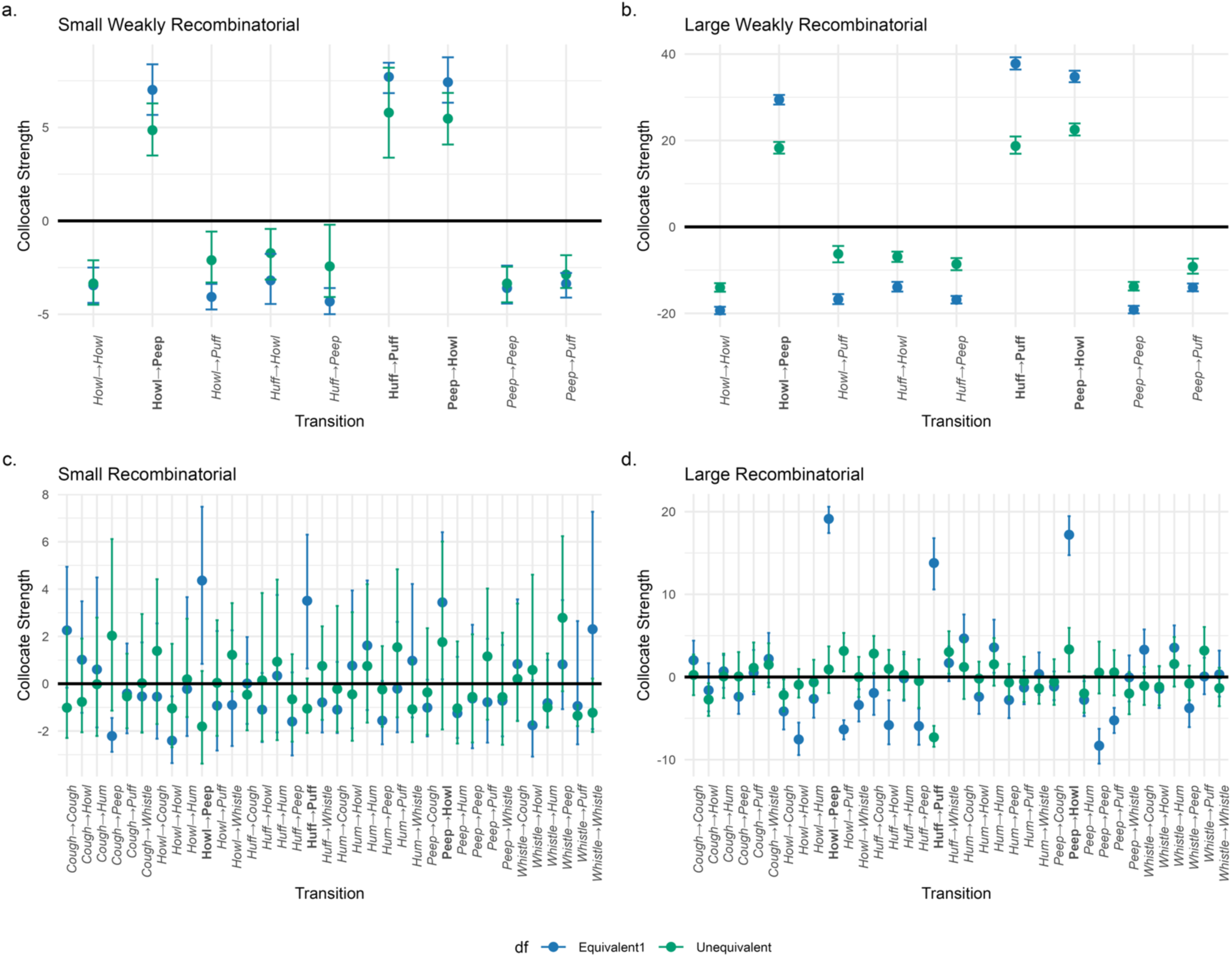
MDCA results comparing *Equivalent* cohort 1 and the *Unequivalent* cohort for each dataset. The top row shows the *Weakly Recombinatorial* data classes, including **a.** *Small* and, **b.** *Large* corpora. The bottom row shows the *Recombinatorial* data classes, including **c.** *Small* and **d.** *Large* corpora. Significant differences between cohorts are identified by no overlap in confidence intervals for a given call bigram type (also see Table 2). Focal bigram types are marked in bold

For all corpora, the frequency distributions for all call bigram types were visualized to ensure our simulated corpora included sampling variation (at both the individual level, and between equivalent cohorts; see Figs. S2-S4). This ensured that our simulations contained some stochastic variation, similarly to when data is sampled from real-world animal populations.

#### 2.2.2 Comparing simulated cohorts

For each dataset, MDCA-Pr was run on all three simulated corpora. We then made two comparisons:

1. A comparison of *Equivalent* cohort 1 and the *Unequivalent* cohort. This comparison allowed us to detect whether imputed differences in call bigram probabilities between cohorts could be detected (a measure of sensitivity).
2. A control comparison between *Equivalent* cohorts 1 and 2. This negative control comparison allowed us to identify whether false-positive differences would be identified between cohorts that had the same underlying probability of producing all call bigram types (a measure of selectivity).

During these comparisons, significant differences in the collocation strength for a given call bigram type were identified by non-overlapping confidence intervals between cohorts (where confidence intervals were Bonferroni corrected to account for the number of pairwise comparisons between cohorts). Overlapping confidence intervals were taken to indicate that there was no significant difference in collocation strength between cohorts for a given bigram type. We compared all bigram types between cohorts, regardless of whether bigrams were identified as significantly attracted in either corpus. This was necessary as for the *Unequivalent* cohort, focal call bigram probabilities were reduced to be very similar to those of non-focal call bigrams. Consequently, all bigram types may re-occur at similar rates in the *Unequivalent* corpus, preventing any subset of bigrams being identified as significantly attracted.

#### 2.2.3 Results

Across all classes of datasets, collocation strength did not significantly differ between *Equivalent* cohorts 1 and 2 for any bigram types, reflecting perfect selectivity of MDCA-Pr during this control intercohort comparison (see Table 2; Fig S5 & Tables S5-S8).

When comparing *Equivalent* cohort 1 with the *Unequivalent* cohort, MDCA-Pr successfully identified differences in collocation strengths for all focal bigram types in both *Large* data classes, and correctly identified a number of differences in the probability of non-focal bigrams occurring in the two corpora (all modified non-focal bigram types for the *Large Weakly Recombinatorial* dataset, and 6/15 modified non-focal bigram types for the *Large Recombinatorial* dataset; see Table 2). Therefore, for *Large* datasets, MDCA-Pr was sensitive to unequal probabilities of calls being combined between cohorts, including consistent detection of differences as low as 5%.

For the *Small* datasets, sensitivity was lower during comparisons of the *Equivalent* cohort 1 and the *Unequivalent* cohort. For the *Small Recombinatorial* dataset, only two focal bigram types were identified as having different collocation strengths between the cohorts (Howl→Peep and Huff→Puff). Whereas, for the *Small Weakly Recombinatorial* dataset, only one focal bigram type was identified differing in collocation strength between cohorts (Huff→Puff). Thus, when datasets are *Small*, MDCA-Pr can be less sensitive to intercohort differences in the probabilities of calls co-occurring, particularly when background recombination is limited.

The sensitivity of our analysis during intercohort comparisons differed considerably between the *Small* and *Large* datasets. As the size of these corpora differ substantially (*Small* = 160 bigrams; *Large* = 3200 bigrams), we ran additional analyses to evaluate whether more moderate increases in sample size would improve sensitivity. Doubling the *Small* sample size (320 bigrams; either by doubling the number of individuals, or the number of calls per individual), improved sensitivity during intercohort comparisons when analysing *Weakly Recombinatorial* data (Double Calls = 2/3 focal bigram types identified as having different collocation strengths; Double Individuals = 3/3; see Fig. S6) whereas sensitivity remained consistent or improved for *Recombinatorial* data (Double Calls = 3/3; Double Individuals = 2/3).

### 2.3 Study 3 – marmoset vocal combinatoriality between sexes

#### 2.3.1 Marmoset vocal data

To provide an example of how MDCA-Pr can be applied to intercohort comparisons of animal vocal data, we investigated whether male and female common marmosets differ in their call combinatoriality when in standardised context, in this case feeding. Common marmosets are a highly vocal primate species, possessing a broad repertoire of call types (Agamaite et al. 2015) that can be flexibly combined into a diverse array of sequences (Bosshard et al. 2024). As cooperative breeders, marmosets engage in active food sharing, and have various short and long-distance calls to signal food availability and individual’s locations to conspecifics (Bosshard et al. 2024; Mircheva et al. 2026). These calls may signal individuals’ own arousal, particularly for females, who get priority access to food (Box et al. 1999; Guerreiro Martins et al. 2019). However, calls may also indicate a general readiness to share food, especially for males, who are more proactive in ensuring food is shared equitably with partners (Mustoe et al. 2016). These discrepancies in food access and sharing between sexes may be reflected within the vocalizations used by either sex when encountering food; for example, males generally produce a higher number of food calls (Mircheva et al. 2026). However, it has not yet been examined whether these behavioural differences between sexes are reflected in the ways in which calls are combined.

We applied MDCA-Pr to an existing corpus of marmoset calls collected by Mircheva et al. (2026). Pairs of monkeys were recorded in two joint 1 x 1.8 x 2 m experimental chambers, where individuals were out of sight from each other (blocked by an opaque screen), but sounds (including vocalizations) could readily travel between either individual in the pair. One monkey of each pair was presented with different stimuli to create distinct behavioural contexts, including an unfamiliar modified rubber duck (inducing an unfamiliar context) or meal worms (inducing a feeding context). Only the individual who was presented with the stimuli was labelled as being in that context. After three minutes, the stimuli were removed, and either a new stimulus was presented to one individual in the pair, or the experiment ended. As data was collected from combinations of known individuals, data collection for this study was not blinded.

To identify the call combinations used by male and female common marmosets when in a feeding context, we sampled all calls produced by a particular individual during the time interval they (and not their partner) were presented with the food stimulus. We restricted our sample to experimental sessions where the pair of individuals were the breeding male and female of each group (2 sessions per pair). This controlled for the effects of developmental stage and breeding status on the dyadic vocal interactions between pairs of marmosets. In total, we initially sampled 10 individuals in pairs from 5 groups.

Our initial sample included 3007 calls (1695 for males; 1312 for females). These calls were part of eight unique call types (see Table S9). Following existing literature on marmoset communication (Meshinska et al. 2024; Bosshard et al. 2024), any calls which occurred within 0.5s of another call were classified as part of the same bout. Additionally, following previous studies of marmoset call combinatoriality, we focused on how marmosets combine different *types* of calls, and therefore condensed any repeats of the same call type within a bout down to a single token in the data (Bosshard et al. 2024). Our analysis of marmoset call-type combinations therefore also reflected Studies 1 & 2, where repeated calls were not included in simulated data. Given repeated calls may be an important aspect of animal combinatoriality (Templeton et al. 2005; Engesser et al. 2017; Berthet et al. 2019), an example applying MDCA-Pr to marmoset call data that included repeats can be found in our supplementary materials. Call-type transitions were sampled from bouts by constructing bigrams that describe all the internal neighbouring call-type combinations. For example, the hypothetical bout ‘*A,B,C,A*’ would be converted into three bigrams: *A→ B; B → C; C → A*.

Whilst this study did not collect new data from Marmoset monkeys, the data used were collected from experiments approved by the Kantonales Veterinäramt Zürich, Switzerland (license number ZH 232/19), and complied with Swiss law (Mircheva et al. 2026).

#### 2.3.2 Applying MDCA-Pr to sex cohorts

MDCA-Pr was applied to the male and female call data. We excluded data for monkeys where fewer than 20 call type bigrams were sampled, representing a trade-off between maximizing the number of individuals included in the data versus the vocal data that could be sampled per individual. This left four males and three females from our initial sample, with a corpus of 733 recorded productions of non-repeating call types, described via 488 pairwise transitions between call types (Males = 266 bigrams; Females = 222) across 246 bouts of vocalization. Note that, as call-type pairs in the same bout are not independent, the total raw sample of call types is not a perfect duplication of pairwise transitions (see the example in the section above that converted four call types into three transitions).

To ensure that the number of males and females sampled for each bootstrap was equal, we sampled with replacement three individuals per sex per bootstrap. To account for uneven numbers of call-type bigrams recorded from different individuals in the data, we modified the bootstrapping procedure to subsample equal amounts of data from each individual monkey, using the smallest sample size for any individual monkey in the data (22 call-type bigrams per individual). Subsampling prevented particularly voluble individuals from being overrepresented in the bootstrapped datasets. Subsampling within individuals was conducted without replacement, following the logic of methodological approaches used to control for unequal cluster sizes in hierarchical or clustered resampling (e.g., Lee et al. 2019). Each bootstrap therefore had a relatively small sample size of 66 call-type bigrams per sex.

For both sexes, we classified significantly attracted call types as those for which the confidence interval for collocate strength was wholly above zero (confidence intervals were adjusted using the same method as Study 2), and where the mean collocation strength for that call-type bigram was above one standard deviation of all mean collocate strengths for a particular sex (as advised from Study 1). Where call types were identified as attracted for both sexes, we evaluated whether collocate strength differed between sexes (where non-overlapping confidence intervals between sexes indicated significant differences in collocate strength; as advised by Study 2). Call-type combinations that were only found in one sex were termed either male-biased or female-biased call-type combinations.

For each attracted combinations of call types, we ran a post-hoc analysis to review which individuals produced each call-type combination in the underlying data. This allowed us to determine whether attracted call types were being produced by many individuals (indicating either species-typical, or sex-biased behaviours), or if they were produced by only one or a few individuals (thus, more likely reflecting idiosyncratic call-type combinations of individuals in either cohort).

#### 2.3.3 Results

We identified a total of five attracted call-type combinations (see Table 3). Two were shared between both sexes: a combination of two long-distance call types (Food Phee→Phee), and a combination of two mobbing call types (Tsik→Ekk). No significant difference in attraction strength was found between the sexes for these shared call-type combinations.

**Table 3.** Attracted call-type combinations identified via MDCA-Pr for male and female marmosets. The mean collocation strength across bootstraps is shown for each call-type combination, alongside the confidence interval for collocation strength. Horizontal dashes indicate that a particular call combination was not significantly attracted for a specific sex. All values rounded to 3.d.p.

| Sex Category | Call combination | Male |  |  | Female |  |  |
| --- | --- | --- | --- | --- | --- | --- | --- |
|  |  | Mean | Lower CI | Upper CI | Mean | Lower CI | Upper CI |
| Both sexes | Tsik→Ekk | 4.481 | 1.600 | 6.068 | 4.677 | 2.415 | 7.140 |
|  | Food Phee→Phee | 3.746 | 2.180 | 5.222 | 3.008 | 1.711 | 6.030 |
| Female-Biased | Ekk→Tsik | - | - | - | 3.900 | 1.919 | 8.000 |
|  | Phee→Short Phee | - | - | - | 4.556 | 3.015 | 6.724 |
|  | Food Peep→Food Phee | - | - | - | 2.307 | 1.539 | 4.264 |

Post-hoc analysis of the underlying data revealed that the Tsik→Ekk call-type combination was produced by all individuals of both sexes, supporting that it is a species-typical behaviour. Post-hoc analysis also identified that the Food Phee→Phee call-type combination was balanced between the sexes in the underlying data; however, it was rare, and produced by one individual of either sex (once by a male, and twice by a female). This combination of call types was likely identified as attracted given that sequences of Food Phees only ever transitioned into sequences of Phees. Despite its rarity, the underlying data supports equal usage by males and females.

We identified three female-biased call-type combinations: Food Peep→Food Phee; Ekk→Tsik; and Phee→Short Phee. The Phee→Short Phee combination was produced by individuals of both sexes. However, within the underlying data, Phee→Short Phee combinations were more commonly produced by females (N_Transitions_ = 16 for males; N_Transitions_ = 24 for females), and males combined Phee calls with a greater diversity of other call types (N_CallsTypes_ = 4 for males, N_CallTypes_ = 3 for females), supporting female-biased call-type attraction. The Food Peep→Food Phee combination was rare, produced once by one female. However, it was the only instance of a Food Phee following another call other than itself. For the female-biased Ekk→Tsik call-type combination, analysis of the data revealed that both sexes produced Ekk→Tsik call-types together (as reflected by the similar mean collocation strength between sexes). However, for males, the confidence interval for this call-type transition overlapped with zero due to a greater uncertainty around true collocate strength (meaning it was not a significant attraction). Whether both Ekk→Tsik and Food Peep→Food Phee represent female-biased call-type combinations would benefit from analysis of a larger dataset.

## 3. Discussion

Collocation analysis has proven to be a useful tool for identifying repeated combinations of signals within animal communication. However current methods for animal collocation analysis lack important uncertainty estimates for collocation strength; overlook the influence of inflated family-wise error induced by multiple comparisons of signal pairs, and ignore nested structure within datasets, all of which reduce the assurance with which animal signal combinations can be validly identified. Additionally, these statistical limitations limit the diversity of questions which can be examined by existing forms of animal collocation analysis, including by preventing cohort-level comparisons of combinatoriality across animals’ full signal repertoires. Using simulated animal-like data, we evaluated the performance of a recently developed form of collocation analysis that can address these statistical limitations (MDCA-Pr), and validate whether it offers a suitable alternative to existing forms of animal collocation analysis. We also evaluated the performance of MDCA-Pr when comparing the combinatorial repertoire of simulated animal cohorts, before providing an example of how this method can be applied to intercohort comparisons of vocal combinatoriality in real animals.

### 3.1 Study 1

Study 1 was used to evaluate the performance of MDCA-Pr on simulated datasets of different sizes and levels of recombinatoriality. Sensitivity of the analysis was generally high across different types of data.

Only one false negative was detected in the *Small Recombinatorial* dataset (the focal bigram type with the lowest combination probability; Huff→Puff, 8%), indicating that MDCA-Pr may only fail to identify call combinations that are relatively weakly attracted if datasets are small and background recombinatoriality is high. The selectivity of MDCA-Pr was also generally high, with false positives only being identified in one dataset (*Large Recombinatorial*). Following a methodological adjustment suggested by van Boekholt et al. (2025), where an additional filter for identifying attraction was implemented based on the standard deviation of all bigrams, our analysis achieved perfect selectivity without any resultant costs to sensitivity. This high selectivity was also reflected in our randomized control data, where no attracted call combinations were detected.

MDCA-Pr therefore performed well on the analysis of animal-like datasets, including those of varying sizes and levels of recombinatoriality. However, our analysis highlights the importance of carefully evaluating the performance of linguistic methods when applied to animal-like data, as additional filters for identifying call attraction were necessary for optimal MDCA performance. Overall, we report that MDCA-Pr offers a useful approach for identifying attracted signal combinations within the communicative repertoires of animals, using methods that possess greater statistical rigour than existing forms of animal collocation analysis. However, we recommend that this method be implemented alongside additional filters that maintains the selectivity with which attracted call combinations are identified (see Fig. 3).

**Figure 3.**
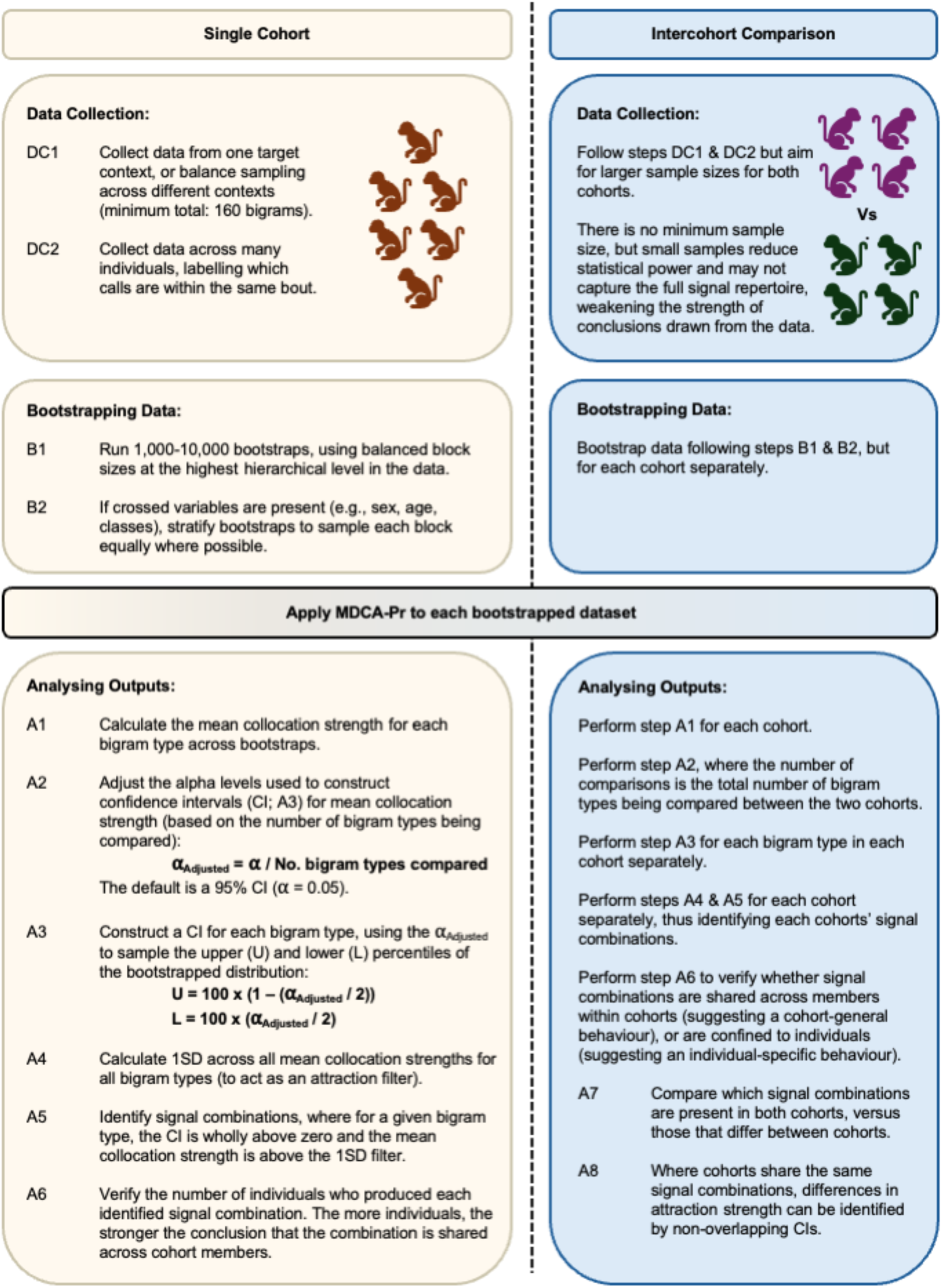
Analytical pipeline for MDCA-Pr. We provide a suggested analytical pipeline for applying MDCA-Pr, followed by further steps to adapt this pipeline for intercohort comparison. R scripts to support implementation of MDCA-Pr can be found in this paper’s supplementary code.

### 3.2 Study 2

In Study 2, we evaluated the performance of MDCA-Pr when identifying differences in the combinatoriality between cohorts of individuals. MDCA-Pr exhibited perfect selectivity during intercohort comparison, with no false-positive differences being identified when corpora were generated using the same probability distribution for bigram types. MDCA-Pr exhibited increasing sensitivity for larger sample sizes. Whilst sensitivity was low for *Small* sample sizes, even moderate increases in sample size (doubling the *Small* sample) markedly improved the analysis’ sensitivity.

Our results validate that MDCA-Pr can detect intercohort differences in signal combinations, and is unlikely to identify false-positive differences between cohorts. However, sample size is an important consideration for ensuring MDCA-Pr is sensitive to more subtle differences between cohorts. Our simulated differences in focal call combination probabilities were relatively small between the *Equivalent* and *Unequivalent* cohorts (between 5-10%; see Table 2), and to detect these differences, several-hundred signal combinations were required. Arguably, meaningful differences between the behaviours of real animal cohorts could be expected to be greater than those we simulated, and in these instances, differences will likely be more easily detected using small samples. Nevertheless, the analysis of small samples should be expected to give conservative estimates of intercohort differences. To estimate minimal sample sizes needed to maintain adequate power of MDCA-Pr during intercohort comparison, researchers could consider simulating their own bespoke datasets of different sizes, tailored to the size of species’ signal repertoire and initial estimates of recombinatoriality. This may then permit evaluation of the minimal sample sizes needed to afford reasonable sensitivity during intercohort comparison, following the protocol we have outlined in Study 2.

### 3.3 Study 3

We followed up Studies 1 & 2 with an example of how MDCA-Pr can be applied to a corpus of vocal data collected from real animals, by comparing the call-type combinations produced by male and female common marmosets in a feeding context, where breeding partners were visually separated from each other and only one dyad partner was presented with food. Our sample size was small for each bootstrap of the data (66 call-type bigrams per sex), therefore the differences we identified are likely conservative (see Study 2). However, three female-biased call-type combinations were identified, as well as two call-type combinations that were shared between sexes (Table 3). Post-hoc analysis revealed that the attracted call-type combinations identified by MDCA-Pr traced how call transitions were distributed across the sexes in the underlying data.

These results suggest that whilst some differences may exist in how male and female marmosets combine call types, these differences are generally small. Whilst behavioural differences in feeding contexts have been identified between the sexes of marmosets (see Section 2.3.1), both sexes engage in food sharing with group members, meaning that within feeding contexts, both sexes may be expected to have largely overlapping communicative goals. Moreover, the motivation to share food varies between individuals, such as through individual-specific variation in hormonal signalling (Finkenwirth et al. 2016), and these individual-level effects may further blur the predictability of sex on individual’s motivation to communicate about food. Before strong biological conclusions are made, an analysis of a larger sample of marmoset vocal data would be beneficial, potentially via combining MDCA-Pr with other analytical tools to determine whether these analyses converge on similar results (e.g. transition probability and positional bias models, see Girard-Buttoz et al. 2022b; though the performance of these methods should also be evaluated when comparing animal cohorts).

Importantly, Study 3 identified further considerations for how MDCA-Pr can be applied rigorously to analyse animal call data. During intersex comparison, post-hoc analyses of underlying data were critical for identifying whether attracted call-type combinations were produced by many individuals within a cohort (thus suggesting a cohort-biased behaviour), or if they were produced by a specific subset of individuals within a cohort (potentially suggesting individual-specific call-type combinations). We anticipate that this additional methodological step is also important for characterizing signal combinatoriality in single populations. Currently, all forms of collocation analysis pool data without consideration of the relative contributions of different individuals within the underlying data. We recommend that in future applications of any collocation analysis, the original data is reviewed to identify whether attracted signal combinations are driven by the behaviours of many individuals, or potentially reflect the behaviour of a small subset of individuals within a sample, and that conclusions are tempered accordingly.

### 3.4. General Discussion

Overall, we report that MDCA-Pr offers an accurate way to identify attracted signal combinations within animal data, and offers a suitable means to compare signal combinatoriality between cohorts of individuals. We provide a recommended analytical pipeline for applying MDCA-Pr to animal signal data, including during cohort comparison (Fig. 3). Within this pipeline, we provide two additional considerations for the collection and analysis of data when using MDCA-Pr:

First, researchers should be mindful of the contexts in which signals are sampled from animals during data collection. In our analysis of marmoset call data, we took a sample of data from a specific context (feeding). We advise that researchers evaluate signal combinatoriality separately across different contexts, as animals may reuse the same signals but combine them in context-specific ways (Berthet et al. 2025; Girard-Buttoz et al. 2025), and this may not be identifiable if data is pooled across contexts. However, if researchers do choose to pool data, we recommend that researchers ensure different contexts are sampled thoroughly and equitably. Similarly, if pooling data across contexts, bootstrapping should be stratified so that each context is consistently represented during error estimation. Stratified bootstraps may also be of use if other variables shared between compared cohorts need to be controlled. For example, when comparing sexes, ensuring that individuals with specific roles, ranks, or age classes are represented.

Second, across our studies we controlled for inflated type 1 error rates during multiple comparison using a Bonferroni correction. Bonferroni corrections can keep type 1 error rates around 5% including up to very high numbers of pairwise comparisons (Bland and Altman 1995). However, these corrections come at the cost of decreasing statistical power with increasing numbers of pairwise comparisons. Thus, when comparing many different bigram types, analyses may be more likely to result in type 2 errors (where valid attraction is not identified within a sample corpus, or a valid difference in attraction is not identified between cohorts). Given our analysis generally showed both high selectivity and sensitivity given reasonable sample sizes, we anticipate Bonferroni corrections will be sufficient in most cases, particularly when making similar numbers of comparisons to those in Studies 1 and 2. However, when working with small samples, or when making many more comparisons, other forms of family-wise error correction could be considered that have less severe consequences for type 2 error rates (e.g. the Holm-Bonferroni method, the Šidák mothod, or Tukey’s method; see Midway et al. 2020).

In conclusion, we demonstrate that MDCA-Pr provides a robust method for identifying attracted signal combinations in animal datasets, overcoming many of the statistical limitations associated with existing approaches to animal collocation analysis. In addition, MDCA-Pr enables direct comparisons of signal attraction across animal cohorts, offering a powerful tool for detecting intraspecific variation in the dynamics of signal combinatoriality within animal communication systems. As with other forms of collocation analysis, MDCA-Pr does not identify the behavioural responses elicited by signal combinations; consequently, attracted signal combinations require further empirical investigation to determine their information content. Nonetheless, as an initial analytical step, the enhanced statistical rigour of MDCA-Pr for identifying signal attraction provides a strong foundation for future examinations of how signal combinatoriality contributes to animal communication, including its potential roles in compositionality (Suzuki et al. 2016; Leroux et al. 2023; Berthet et al. 2025) and pragmatic inference (Schlenker et al. 2016a).

## Acknowledgements

This research was funded by the NCCR Evolving Language (Swiss NSF agreement Nr. 51NF40_180888) and the European Research Council (ERC) under the European Union’s Horizon 2020 research and innovation programme (grant agreement No. 101001295). We would like to thank Stefan Gries and Alexandra Bosshard, whose previously published work has informed this research project.

## Data & Code availability

Data and code associated with this manuscript can be found at https://doi.org/10.17605/OSF.IO/6CGBM

## Notes

https://doi.org/10.17605/OSF.IO/6CGBM

### Competing Interest Statement

The authors have declared no competing interest.

### Summary of Updates

We streamlined the title to highlight the major contributions of this manuscript.

## References

Agamaite JA, Chang C-J, Osmanski MS, Wang X (2015) A quantitative acoustic analysis of the vocal repertoire of the common marmoset (*Callithrix jacchus*). J. Acoust. Soc. Am. 138:2906–2928. 10.1121/1.4934268

Bartsch S (2004) Structural and Functional Properties of Collocations in English: A corpus study of lexical and pragmatic constraints on lexical co-occurence. Gunter Narr Verlag, Tübingen

Berthet M, Mesbahi G, Pajot A, et al (2019) Titi monkeys combine alarm calls to create probabilistic meaning. Sci. Adv. 5:eaav3991. 10.1126/sciadv.aav3991

Berthet M, Surbeck M, Townsend SW (2025) Extensive compositionality in the vocal system of bonobos. Science 388:104–108. 10.1126/science.adv1170

Bland JMartin, Altman DG (1995) Multiple significance tests: the Bonferroni method. BMJ 310:170

Bosshard AB, Burkart JM, Merlo P, et al (2024) Beyond bigrams: call sequencing in the common marmoset (*Callithrix jacchus*) vocal system. R. Soc. Open Sci. 11:240218. 10.1098/rsos.240218

Bosshard AB, Leroux M, Lester NA, et al (2022) From collocations to call-locations : using linguistic methods to quantify animal call combinations. Behav. Ecol. Sociobiol. 76:1–8. 10.1007/s00265-022-03224-3

Box H, Yamamoto ME, Lopes FA (1999) Gender Differences in Marmosets and Tamarins: Responses to Food Tasks. Int. J. Comp. Psychol. 12:59–70. 10.46867/C42G67

Briseño-Jaramillo M, Sosa-López JR, Ramos-Fernández G, Lemasson A (2022) Flexible use of contact calls in a species with high fission–fusion dynamics. Phil. Trans. R. Soc. B 377:20210309. 10.1098/rstb.2021.0309

Church K, Gale W, Hanks P, Hindle D (1991) Using Statistics in Lexical Analysis. In: Zernik U (ed) Lexical Acquisition: Exploiting On-Line Resources to Build a Lexicon, 1st edn. Psychology Press, New York, NY, pp 115–164

Collier K, Bickel B, Van Schaik CP, et al (2014) Language evolution: syntax before phonology? Proc. R. Soc. B 281:20140263. 10.1098/rspb.2014.0263

Collier K, Radford AN, Stoll S, et al (2020) Dwarf mongoose alarm calls: investigating a complex non-human animal call. Proc. R. Soc. B 287:20192514. 10.1098/rspb.2019.2514

Collier K, Townsend SW, Manser MB (2017) Call concatenation in wild meerkats. Anim. Behav. 134:257–269. 10.1016/j.anbehav.2016.12.014

Coye C, Ouattara K, Arlet ME, et al (2018a) Flexible use of simple and combined calls in female Campbell’s monkeys. Anim Behav 141:171–181. 10.1016/j.anbehav.2018.05.014

Coye C, Townsend S, Lemasson A (2018b) From animal communication to linguistics and back: Insight from combinatorial abilities in monkeys and birds. In: Boë L-J, Fagot J, Perrier P, Schwartz J-L (eds) Origins of Human Language: Continuities and Discontinuities with Nonhuman Primates. Peter Lang GmbH, Berlin

Demartsev V, Averly B, Johnson-Ulrich L, et al (2024) Mapping vocal interactions in space and time differentiates signal broadcast versus signal exchange in meerkat groups. Phil. Trans. R. Soc. B 379:20230188. 10.1098/rstb.2023.0188

Engesser S, Holub JL, O’Neill LG, et al (2019) Chestnut-crowned babbler calls are composed of meaningless shared building blocks. Proc. Nat. Acad. Sci. U.S.A. 116:19579–19584. 10.1073/pnas.1819513116

Engesser S, Ridley AR, Townsend SW (2016) Meaningful call combinations and compositional processing in the southern pied babbler. Proc. Nat. Acad. Sci. U.S.A. 113:5976–5981. 10.1073/pnas.1600970113

Engesser S, Ridley AR, Townsend SW (2017) Element repetition rates encode functionally distinct information in pied babbler ‘clucks’ and ‘purrs.’ Anim. Cogn. 20:953–960. 10.1007/s10071-017-1114-6

Engesser S, Ridley AR, Watson SK, et al (2024) Seeds of language-like generativity in bird call combinations. Proc. R. Soc. B 291:20240922. 10.1098/rspb.2024.0922

Engesser S, Townsend SW (2019) Combinatoriality in the vocal systems of nonhuman animals. WIREs Cogn. Sci. 10:e1493. 10.1002/wcs.1493

Evert S (2009) Corpora and collocations. In: Lüdeling A, Kytö M (eds) Corpus Linguistics. An International Handbook. Mouton de Gruyter, pp 1212–1248

Finkenwirth C, Martins E, Deschner T, Burkart JM (2016) Oxytocin is associated with infant-care behavior and motivation in cooperatively breeding marmoset monkeys. Horm. Behav. 80:10–18. 10.1016/j.yhbeh.2016.01.008

Fitch WT (2018) What animals can teach us about human language: the phonological continuity hypothesis. Curr. Opin. Behav. Sci. 21:68–75. 10.1016/j.cobeha.2018.01.014

Freymann E, d’Oliveira Coelho J, Hobaiter C, et al (2024) Applying collocation and APRIORI analyses to chimpanzee diets: Methods for investigating nonrandom food combinations in primate self-medication. Am. J. Primatol. 86:e23603. 10.1002/ajp.23603

Girard-Buttoz C, Bortolato T, Laporte M, et al (2022a) Population-specific call order in chimpanzee greeting vocal sequences. iScience 25:104851. 10.1016/j.isci.2022.104851

Girard-Buttoz C, Neumann C, Bortolato T, et al (2025) Versatile use of chimpanzee call combinations promotes meaning expansion. Sci. Adv. 11:eadq2879. 10.1126/sciadv.adq2879

Girard-Buttoz C, Zaccarella E, Bortolato T, et al (2022b) Chimpanzees produce diverse vocal sequences with ordered and recombinatorial properties. Commun. Biol. 5:410. 10.1038/s42003-022-03350-8

Gries S (2024) Collostructional analysis: Computing the degree of association between words and words/constructions v4.1. https://www.stgries.info/teaching/groningen/coll.analysis.r accessed 10th August 2025

Gries STh (2022) Toward more careful corpus statistics: uncertainty estimates for frequencies, dispersions, association measures, and more. RMAL 1:100002. 10.1016/j.rmal.2021.100002

Gries STh (2023) Overhauling Collostructional Analysis: Towards More Descriptive Simplicity and More Explanatory Adequacy. Cogn. Semant. 9:351–386. 10.1163/23526416-bja10056

Gu H, Sun C, Gong L, et al (2023) Sex ratio potentially influence the complexity of social calls in Himalayan leaf-nosed bat groups. Front. Ecol. Evol. 11:955540. 10.3389/fevo.2023.955540

Guerreiro Martins EM, Moura AC d. A, Finkenwirth C, et al (2019) Food sharing patterns in three species of callitrichid monkeys (*Callithrix jacchus, Leontopithecus chrysomelas, Saguinus midas*): Individual and species differences. J. Comp. Psychol. 133:474–487. 10.1037/com0000169

Hedwig D, Kohlberg A (2024) Call combination in African forest elephants *Loxodonta cyclotis*. PLoS ONE 19:e0299656. 10.1371/journal.pone.0299656

Idsardi WJ (2019) Some cautions regarding the phonological continuity hypothesis. Phil. Trans. R. Soc. B 375:20190050. 10.1098/rstb.2019.0050

Le Floch A, Girard-Buttoz C, Azaiez TS, et al (2026) Rule-based sequences in sooty mangabey vocal communication. R. Soc. Open. Sci. 13(2):251944. 10.1098/rsos.251944

Lee D, Kim J, Skinner C (2019) Within-cluster resampling for multilevel models under informative cluster size. Biometrika 106:965–972. 10.1093/biomet/asz035

Lehecka T (2015) Collocation and colligation. In: Handbook of Pragmatics. John Benjamins Publishing Company, pp 1–20. 10.1075/hop.19.col2

Leroux M, Bosshard AB, Chandia B, et al (2021) Chimpanzees combine pant hoots with food calls into larger structures. Anim. Behav. 179:41–50. 10.1016/j.anbehav.2021.06.026

Leroux M, Chandia B, Bosshard AB, et al (2022) Call combinations in chimpanzees: a social tool? Behav. Ecol. 33:1036–1043. 10.1093/beheco/arac074

Leroux M, Schel, Anne M, Wilke, Claudia, et al (2023) Call combinations and compositional processing in wild chimpanzees. Nat. Commun. 14:1–8. 10.1038/s41467-023-37816-y

Leroux M, Townsend SW (2020) Call combinations in great apes and the evolution of syntax. AB&C 7:131–139. 10.26451/abc.07.02.07.2020

Liao S, Gries STh, Wulff S (2024) Transfer five ways: applications of multiple distinctive collexeme analysis to the dative alternation in Mandarin Chinese. CLLT. 10.1515/cllt-2024-0033

Meshinska K, Burkart JM, Bell MBV, Wierucka K (2024) Echoes of self: Understanding acoustic structure and informational content in common marmoset (*Callithrix jacchus*) phee sequences. bioRxiv. 10.1101/2024.04.14.589400

Midway S, Robertson M, Flinn S, Kaller M (2020) Comparing multiple comparisons: practical guidance for choosing the best multiple comparisons test. PeerJ 8:e10387. 10.7717/peerj.10387

Mine JG, Dees LC, Wilke C, et al (2025) Chimpanzee mothers, but not fathers, influence offspring vocal–visual communicative behavior. PLoS Biol. 23:e3003270. 10.1371/journal.pbio.3003270

Mine JG, Wilke C, Zulberti C, et al (2024) Vocal-visual combinations in wild chimpanzees. Behav. Ecol. Sociobiol. 78:108. 10.1007/s00265-024-03523-x

Mircheva M, Brügger R, Burkart JM (2026) Who says what when? Patterns in captive common marmoset (Callithrix jacchus) volubility. 10.64898/2026.01.24.701497

Mustoe AC, Harnisch AM, Hochfelder B, et al (2016) Inequity aversion strategies between marmosets are influenced by partner familiarity and sex but not by oxytocin. Anim. Behav. 114:69–79. 10.1016/j.anbehav.2016.01.025

Nesselhauf N (2005) Collocations in a Learner Corpus. John Benjamins B.V., Amsterdam, The Netherlands

Nowak MA, Plotkin JB, Jansen VAA (2000) The evolution of syntactic communication. Nature 404:495–498. 10.1038/35006635

R Core Team (2022) R: A language and environment for statistical computing. https://www.R-project.org/

Schamberg I, Surbeck M, Townsend SW (2024) Cross-population variation in usage of a call combination: evidence of signal usage flexibility in wild bonobos. Anim. Cogn. 27:58. 10.1007/s10071-024-01884-4

Schlenker P, Chemla E, Schel AM, et al (2016a) Formal monkey linguistics. Theor. Linguist. 42:1–90. 10.1515/tl-2016-0001

Schlenker P, Chemla E, Schel AM, et al (2016b) Formal monkey linguistics: The debate. Theor. Linguist. 42:173–201. 10.1515/tl-2016-0010

Schlenker P, Chlema E, Arnold K, et al (2014) Monkey semantics: two “dialects” of Campbell’s monkey alarm calls. Linguist. Philos. 37:439–501. 10.1007/s

Schlenker P, Salis A, Leroux M, et al (2024) Minimal Compositionality *versus* Bird Implicatures: two theories of ABC-D sequences in Japanese tits. Biol. Rev. brv.13068. 10.1111/brv.13068

Spiess S, Mylne HK, Engesser S, et al (2022) Syntax-like Structures in Maternal Contact Calls of Chestnut-Crowned Babblers (*Pomatostomus ruficeps*). Int. J. Primatol. 10.1007/s10764-022-00332-9

Stefanowitsch A, Gries STh (2003) Collostructions: Investigating the interaction of words and constructions. IJCL 8:209–243. 10.1075/ijcl.8.2.03ste

Stefanowitsch A, Gries STh (2005) Covarying collexemes. CLLT 1:1–43. 10.1515/cllt.2005.1.1.1

Suzuki TN, Wheatcroft D, Griesser M (2016) Experimental evidence for compositional syntax in bird calls. Nat. Commun. 7:10986. 10.1038/ncomms10986

Suzuki TN, Zuberbühler K (2019) Animal syntax. Curr. Biol. 29:R669–R671. 10.1016/j.cub.2019.05.045

Templeton CN, Greene E, Davis K (2005) Allometry of Alarm Calls: Black-Capped Chickadees Encode Information About Predator Size. Science 308:1934–1937. 10.1126/science.1108841

Townsend SW, Engesser S, Stoll S, et al (2018) Compositionality in animals and humans. PLoS Biol. 16:e2006425. 10.1371/journal.pbio.2006425

van Boekholt B, Bosshard AB, Pika S (2025) Sequence organization of mother–infant interactions in chimpanzees *(Pan troglodytes schweinfurthii)* in the wild. Proc. R. Soc. B. 292:20252271. 10.1098/rspb.2025.2271

Walsh SL, Townsend SW, Engesser S, Ridley AR (2024) Call combination production is linked to the social environment in Western Australian magpies (*Gymnorhina tibicen dorsalis*). Phil. Trans. R. Soc. B. 379:20230198. 10.1098/rstb.2023.0198

Wewhare N, Krishnan A (2024) Individual-specific associations between warble song notes and body movements in budgerigar courtship displays. BiO 13:bio060497. 10.1242/bio.060497

Xiao R, McEnery T (2006) Collocation, Semantic Prosody, and Near Synonymy: A Cross-Linguistic Perspective. Appl. Linguist. 27:103–129. 10.1093/applin/ami045

Zuberbühler K (2019) Syntax and compositionality in animal communication. Phil. Trans. R. Soc. B. 375:20190062. 10.1098/rstb.2019.0062

