## Supplementary Materials for "Animal collocation analysis 2.0: improving statistical inference and applications for cohort comparisons"

Contents:

| | |
|---|---|
| Table S1 | Page 2 |
| Table S2 | Page 2 |
| Table S3 | Page 3 |
| Table S4 | Page 4 |
| Figure S1 | Page 5 |
| Figure S2 | Page 6 |
| Figure S3 | Page 7 |
| Figure S4 | Page 8 |
| Figure S5 | Page 9 |
| Table S5 | Page 10 |
| Table S6 | Page 11 |
| Table S7 | Page 12 |
| Table S8 | Page 14 |
| Figure S6 | Page 16 |
| Table S9 | Page 17 |
| Table S10 | Page 18 |
| Table S11 | Page 19 |
| Table S12 | Page 20 |
| Marmoset results with repeating calls | Page 20 |

**Table S1. MDCA-Pr results for the *Small Exclusive* dataset.** All values rounded to 3.d.p. AttRep indicates attraction and repulsion before applying the 1SD filter.

| call | collocate | samples | means | upper | lower | encompass_zero | significantToZero | Direction | SD |
|---|---|---|---|---|---|---|---|---|---|
| Howl | Howl | 10000 | -4.706 | -4.137 | -5.200 | FALSE | TRUE | Repulsion | 6.316 |
| Howl | Peep | 10000 | 7.116 | 8.300 | 5.930 | FALSE | TRUE | Attraction | 6.316 |
| Howl | Puff | 10000 | -4.116 | -3.500 | -4.773 | FALSE | TRUE | Repulsion | 6.316 |
| Huff | Howl | 10000 | -3.510 | -2.943 | -4.104 | FALSE | TRUE | Repulsion | 6.316 |
| Huff | Peep | 10000 | -4.116 | -3.500 | -4.773 | FALSE | TRUE | Repulsion | 6.316 |
| Huff | Puff | 10000 | 9.570 | 10.436 | 8.696 | FALSE | TRUE | Attraction | 6.316 |
| Peep | Howl | 10000 | 8.613 | 9.961 | 7.431 | FALSE | TRUE | Attraction | 6.316 |
| Peep | Peep | 10000 | -4.706 | -4.137 | -5.196 | FALSE | TRUE | Repulsion | 6.316 |
| Peep | Puff | 10000 | -3.510 | -2.943 | -4.104 | FALSE | TRUE | Repulsion | 6.316 |

**Table S2. MDCA-Pr results for the *Large Exclusive* dataset.** All values rounded to 3.d.p. AttRep indicates attraction and repulsion before applying the 1SD filter.

| call | collocate | samples | means | upper | lower | encompass_zero | significantToZero | Direction | SD |
|---|---|---|---|---|---|---|---|---|---|
| Howl | Howl | 10000 | -20.901 | -20.388 | -21.378 | FALSE | TRUE | Repulsion | 28.281 |
| Howl | Peep | 10000 | 31.378 | 32.456 | 30.264 | FALSE | TRUE | Attraction | 28.281 |
| Howl | Puff | 10000 | -18.797 | -18.332 | -19.293 | FALSE | TRUE | Repulsion | 28.281 |
| Huff | Howl | 10000 | -15.599 | -15.013 | -16.169 | FALSE | TRUE | Repulsion | 28.281 |
| Huff | Peep | 10000 | -18.797 | -18.332 | -19.293 | FALSE | TRUE | Repulsion | 28.281 |
| Huff | Puff | 10000 | 42.536 | 43.558 | 41.472 | FALSE | TRUE | Attraction | 28.281 |
| Peep | Howl | 10000 | 39.223 | 40.005 | 38.467 | FALSE | TRUE | Attraction | 28.281 |
| Peep | Peep | 10000 | -20.901 | -20.388 | -21.378 | FALSE | TRUE | Repulsion | 28.281 |
| Peep | Puff | 10000 | -15.599 | -15.013 | -16.169 | FALSE | TRUE | Repulsion | 28.281 |

**Table S3. MDCA-Pr results for the *Small Recombinatorial* dataset.** All values rounded to 3.d.p. AttRep indicates attraction and repulsion before applying the 1SD filter.

| call | collocate | samples | means | upper | lower | encompass_zero | significantToZero | Direction | SD |
|---|---|---|---|---|---|---|---|---|---|
| Cough | Cough | 10000 | -0.316 | 2.477 | -1.759 | TRUE | FALSE | NA | 1.554 |
| Cough | Howl | 10000 | 0.645 | 3.282 | -1.919 | TRUE | FALSE | NA | 1.554 |
| Cough | Hum | 10000 | 1.074 | 3.855 | -1.561 | TRUE | FALSE | NA | 1.554 |
| Cough | Peep | 10000 | -1.402 | 0.853 | -3.000 | TRUE | FALSE | NA | 1.554 |
| Cough | Puff | 10000 | 0.328 | 2.195 | -1.944 | TRUE | FALSE | NA | 1.554 |
| Cough | Whistle | 10000 | 0.037 | 2.783 | -1.997 | TRUE | FALSE | NA | 1.554 |
| Howl | Cough | 10000 | -0.914 | 1.732 | -2.485 | TRUE | FALSE | NA | 1.554 |
| Howl | Howl | 10000 | -1.812 | 0.545 | -3.228 | TRUE | FALSE | NA | 1.554 |
| Howl | Hum | 10000 | -0.303 | 2.483 | -1.917 | TRUE | FALSE | NA | 1.554 |
| Howl | Peep | 10000 | 3.842 | 5.126 | 2.750 | FALSE | TRUE | Attraction | 1.554 |
| Howl | Puff | 10000 | -0.303 | 1.909 | -2.337 | TRUE | FALSE | NA | 1.554 |
| Howl | Whistle | 10000 | -1.602 | 0.938 | -3.305 | TRUE | FALSE | NA | 1.554 |
| Huff | Cough | 10000 | -0.810 | 1.932 | -1.984 | TRUE | FALSE | NA | 1.554 |
| Huff | Howl | 10000 | -1.914 | 0.390 | -3.146 | TRUE | FALSE | NA | 1.554 |
| Huff | Hum | 10000 | 0.738 | 2.137 | -0.901 | TRUE | FALSE | NA | 1.554 |
| Huff | Peep | 10000 | -1.491 | 0.136 | -2.969 | TRUE | FALSE | NA | 1.554 |
| Huff | Puff | 10000 | 1.947 | 3.895 | -0.828 | TRUE | FALSE | NA | 1.554 |
| Huff | Whistle | 10000 | 2.140 | 4.942 | -1.369 | TRUE | FALSE | NA | 1.554 |
| Hum | Cough | 10000 | 1.732 | 5.070 | -1.232 | TRUE | FALSE | NA | 1.554 |
| Hum | Howl | 10000 | -1.105 | 1.076 | -2.129 | TRUE | FALSE | NA | 1.554 |
| Hum | Hum | 10000 | 0.486 | 4.107 | -1.658 | TRUE | FALSE | NA | 1.554 |
| Hum | Peep | 10000 | -0.230 | 1.854 | -2.095 | TRUE | FALSE | NA | 1.554 |
| Hum | Puff | 10000 | -0.266 | 2.743 | -2.345 | TRUE | FALSE | NA | 1.554 |
| Hum | Whistle | 10000 | 0.224 | 4.410 | -2.009 | TRUE | FALSE | NA | 1.554 |
| Peep | Cough | 10000 | -1.344 | -0.548 | -2.178 | FALSE | TRUE | Repulsion | 1.554 |
| Peep | Howl | 10000 | 4.808 | 7.487 | 1.414 | FALSE | TRUE | Attraction | 1.554 |
| Peep | Hum | 10000 | -0.752 | 2.240 | -1.896 | TRUE | FALSE | NA | 1.554 |
| Peep | Peep | 10000 | -1.082 | 0.653 | -2.598 | TRUE | FALSE | NA | 1.554 |
| Peep | Puff | 10000 | -1.931 | -0.848 | -3.000 | FALSE | TRUE | Repulsion | 1.554 |
| Peep | Whistle | 10000 | -0.529 | 1.311 | -1.837 | TRUE | FALSE | NA | 1.554 |
| Whistle | Cough | 10000 | 2.268 | 5.774 | -1.255 | TRUE | FALSE | NA | 1.554 |
| Whistle | Howl | 10000 | 0.409 | 2.922 | -2.122 | TRUE | FALSE | NA | 1.554 |
| Whistle | Hum | 10000 | -1.417 | -0.474 | -2.305 | FALSE | TRUE | Repulsion | 1.554 |
| Whistle | Peep | 10000 | -1.023 | 0.479 | -2.174 | TRUE | FALSE | NA | 1.554 |
| Whistle | Puff | 10000 | 0.105 | 2.691 | -2.081 | TRUE | FALSE | NA | 1.554 |
| Whistle | Whistle | 10000 | 0.226 | 3.653 | -2.054 | TRUE | FALSE | NA | 1.554 |

**Table S4. MDCA-Pr results for the *Large Recombinatorial* dataset.** All values rounded to 3.d.p. AttRep indicates attraction and repulsion before applying the 1SD filter.

| call | collocate | samples | means | upper | lower | encompass_zero | significantToZero | AttRep | SD |
|---|---|---|---|---|---|---|---|---|---|
| Cough | Cough | 10000 | 4.474 | 7.346 | 1.726 | FALSE | TRUE | Attraction | 5.989 |
| Cough | Howl | 10000 | -2.170 | -0.128 | -4.323 | FALSE | TRUE | Repulsion | 5.989 |
| Cough | Hum | 10000 | 2.287 | 5.898 | -0.716 | TRUE | FALSE | NA | 5.989 |
| Cough | Peep | 10000 | -4.025 | -1.617 | -6.356 | FALSE | TRUE | Repulsion | 5.989 |
| Cough | Puff | 10000 | -1.690 | 0.740 | -3.921 | TRUE | FALSE | NA | 5.989 |
| Cough | Whistle | 10000 | 3.813 | 7.670 | 0.782 | FALSE | TRUE | Attraction | 5.989 |
| Howl | Cough | 10000 | -3.962 | -1.559 | -6.123 | FALSE | TRUE | Repulsion | 5.989 |
| Howl | Howl | 10000 | -6.216 | -3.588 | -8.601 | FALSE | TRUE | Repulsion | 5.989 |
| Howl | Hum | 10000 | -3.719 | -1.534 | -5.815 | FALSE | TRUE | Repulsion | 5.989 |
| Howl | Peep | 10000 | 18.493 | 20.300 | 16.730 | FALSE | TRUE | Attraction | 5.989 |
| Howl | Puff | 10000 | -6.527 | -4.093 | -8.518 | FALSE | TRUE | Repulsion | 5.989 |
| Howl | Whistle | 10000 | -2.968 | -1.239 | -4.770 | FALSE | TRUE | Repulsion | 5.989 |
| Huff | Cough | 10000 | -0.614 | 1.908 | -2.906 | TRUE | FALSE | NA | 5.989 |
| Huff | Howl | 10000 | -4.189 | -2.419 | -5.858 | FALSE | TRUE | Repulsion | 5.989 |
| Huff | Hum | 10000 | -1.229 | 1.323 | -3.932 | TRUE | FALSE | NA | 5.989 |
| Huff | Peep | 10000 | -6.611 | -4.741 | -8.402 | FALSE | TRUE | Repulsion | 5.989 |
| Huff | Puff | 10000 | 14.834 | 16.739 | 12.471 | FALSE | TRUE | Attraction | 5.989 |
| Huff | Whistle | 10000 | -1.325 | 1.585 | -4.245 | TRUE | FALSE | NA | 5.989 |
| Hum | Cough | 10000 | 2.686 | 5.973 | -0.173 | TRUE | FALSE | NA | 5.989 |
| Hum | Howl | 10000 | -3.481 | -1.693 | -5.169 | FALSE | TRUE | Repulsion | 5.989 |
| Hum | Hum | 10000 | 4.001 | 7.182 | 1.256 | FALSE | TRUE | Attraction | 5.989 |
| Hum | Peep | 10000 | -3.206 | -0.823 | -5.065 | FALSE | TRUE | Repulsion | 5.989 |
| Hum | Puff | 10000 | -0.471 | 1.955 | -3.025 | TRUE | FALSE | NA | 5.989 |
| Hum | Whistle | 10000 | 2.834 | 5.956 | -0.267 | TRUE | FALSE | NA | 5.989 |
| Peep | Cough | 10000 | -1.879 | 0.445 | -4.149 | TRUE | FALSE | NA | 5.989 |
| Peep | Howl | 10000 | 16.866 | 19.644 | 13.804 | FALSE | TRUE | Attraction | 5.989 |
| Peep | Hum | 10000 | -1.806 | 0.616 | -4.241 | TRUE | FALSE | NA | 5.989 |
| Peep | Peep | 10000 | -6.504 | -4.685 | -8.249 | FALSE | TRUE | Repulsion | 5.989 |
| Peep | Puff | 10000 | -5.007 | -3.722 | -6.549 | FALSE | TRUE | Repulsion | 5.989 |
| Peep | Whistle | 10000 | -2.314 | -0.174 | -4.452 | FALSE | TRUE | Repulsion | 5.989 |
| Whistle | Cough | 10000 | 1.646 | 4.663 | -1.112 | TRUE | FALSE | NA | 5.989 |
| Whistle | Howl | 10000 | -2.288 | 0.076 | -4.637 | TRUE | FALSE | NA | 5.989 |
| Whistle | Hum | 10000 | 2.927 | 4.938 | 1.036 | FALSE | TRUE | Attraction | 5.989 |
| Whistle | Peep | 10000 | -2.446 | -0.194 | -4.442 | FALSE | TRUE | Repulsion | 5.989 |
| Whistle | Puff | 10000 | -0.372 | 1.941 | -2.312 | TRUE | FALSE | NA | 5.989 |
| Whistle | Whistle | 10000 | 2.242 | 6.440 | -1.417 | TRUE | FALSE | NA | 5.989 |

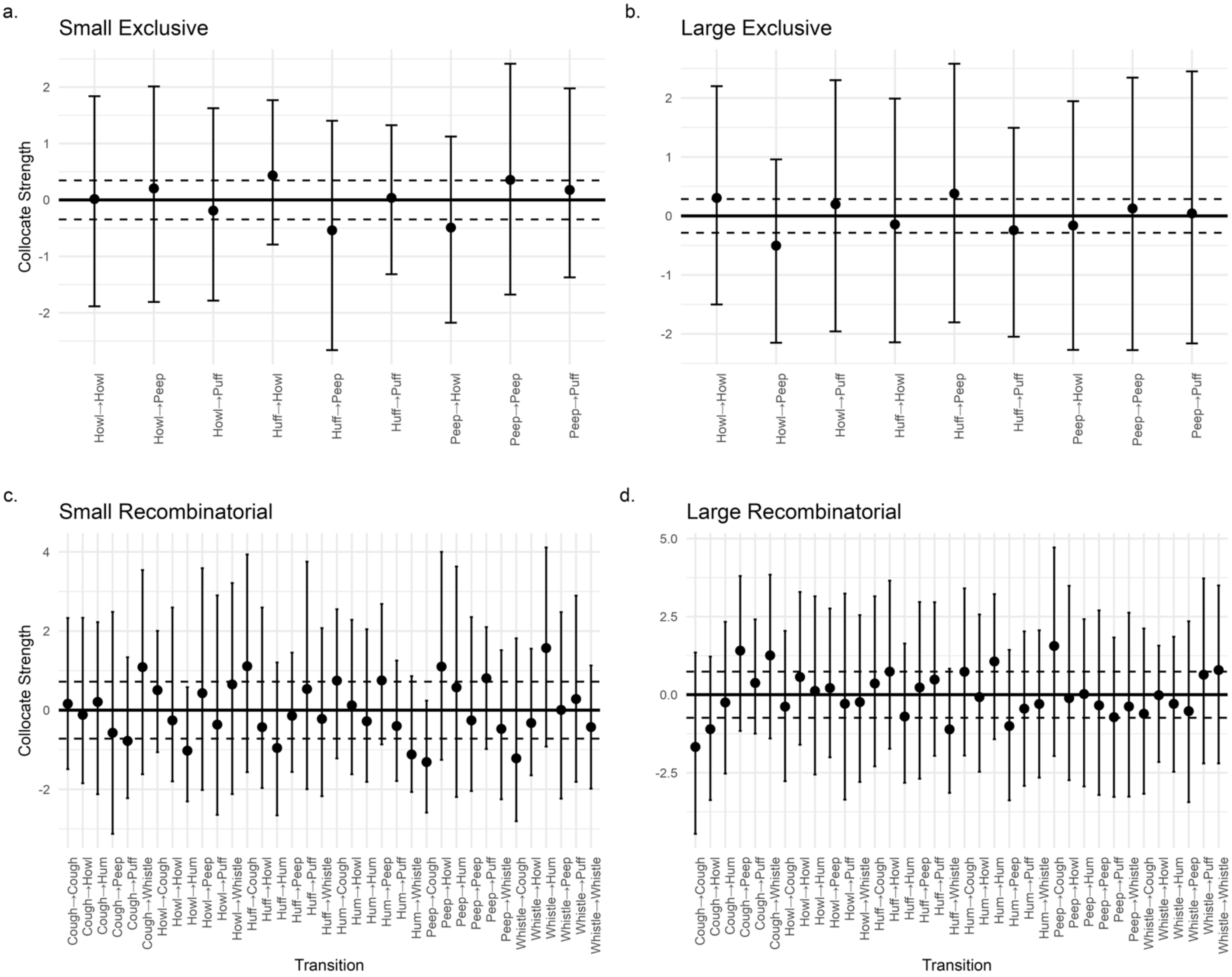

**Fig S1. Output of MDCA-Pr on four datasets with a uniform random distribution of bigram types.** A comparable randomized dataset was generated for each dataset in Study 1 as a negative control: **a.** the *Small Exclusive* dataset; **b.** the *Large Exclusive* dataset; **c.** the *Small Recombinatorial* dataset, and **d.** the *Large Recombinatorial* dataset. No significant attraction was detected, as confidence intervals overlapped with 0 for all bigram types across all datasets.

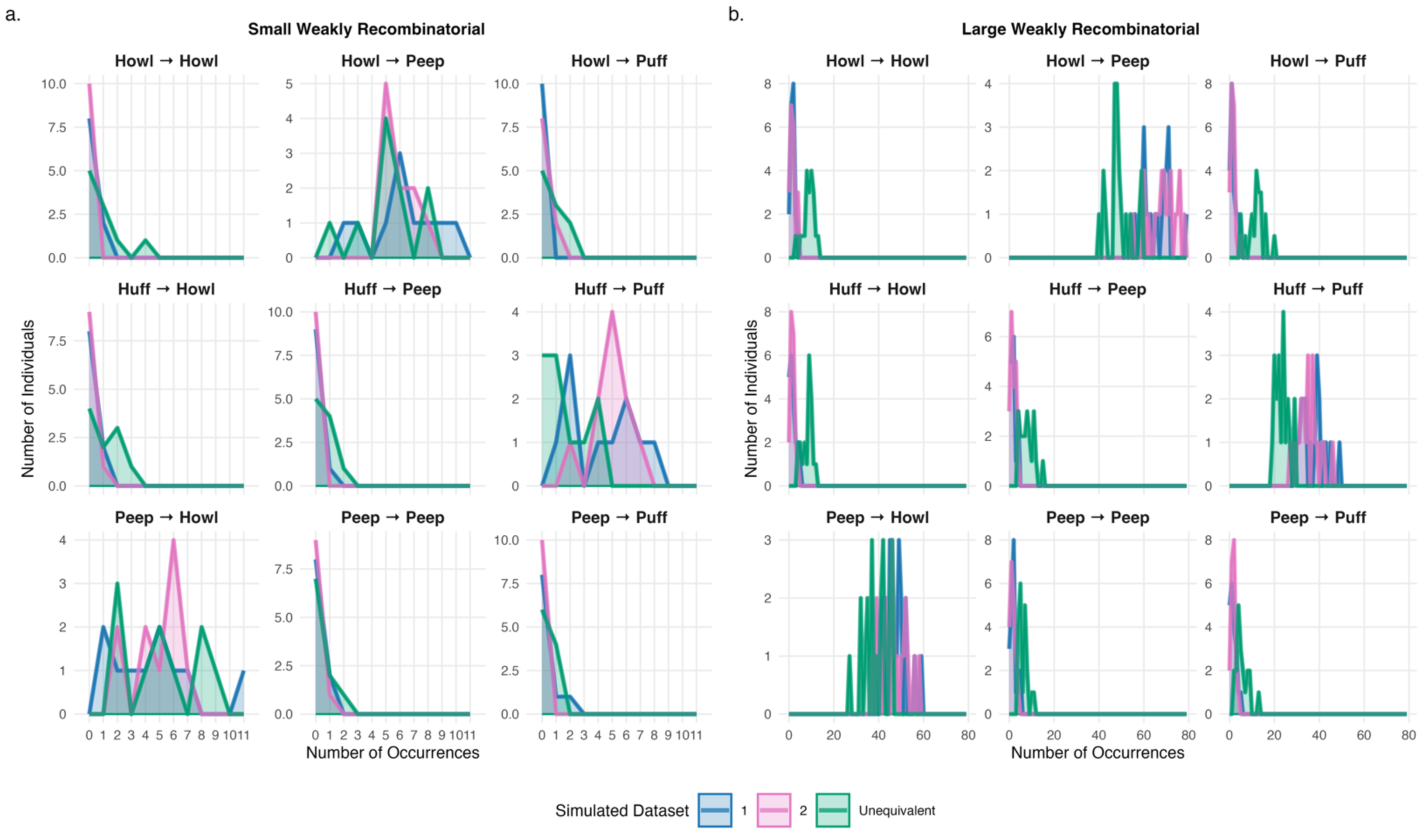

**Fig. S2 Frequency distributions for bigram types per simulated individual across the *Weakly Recombinatorial* corpora from Study 2. a.** The *Small* dataset. **b.** The *Large* dataset.

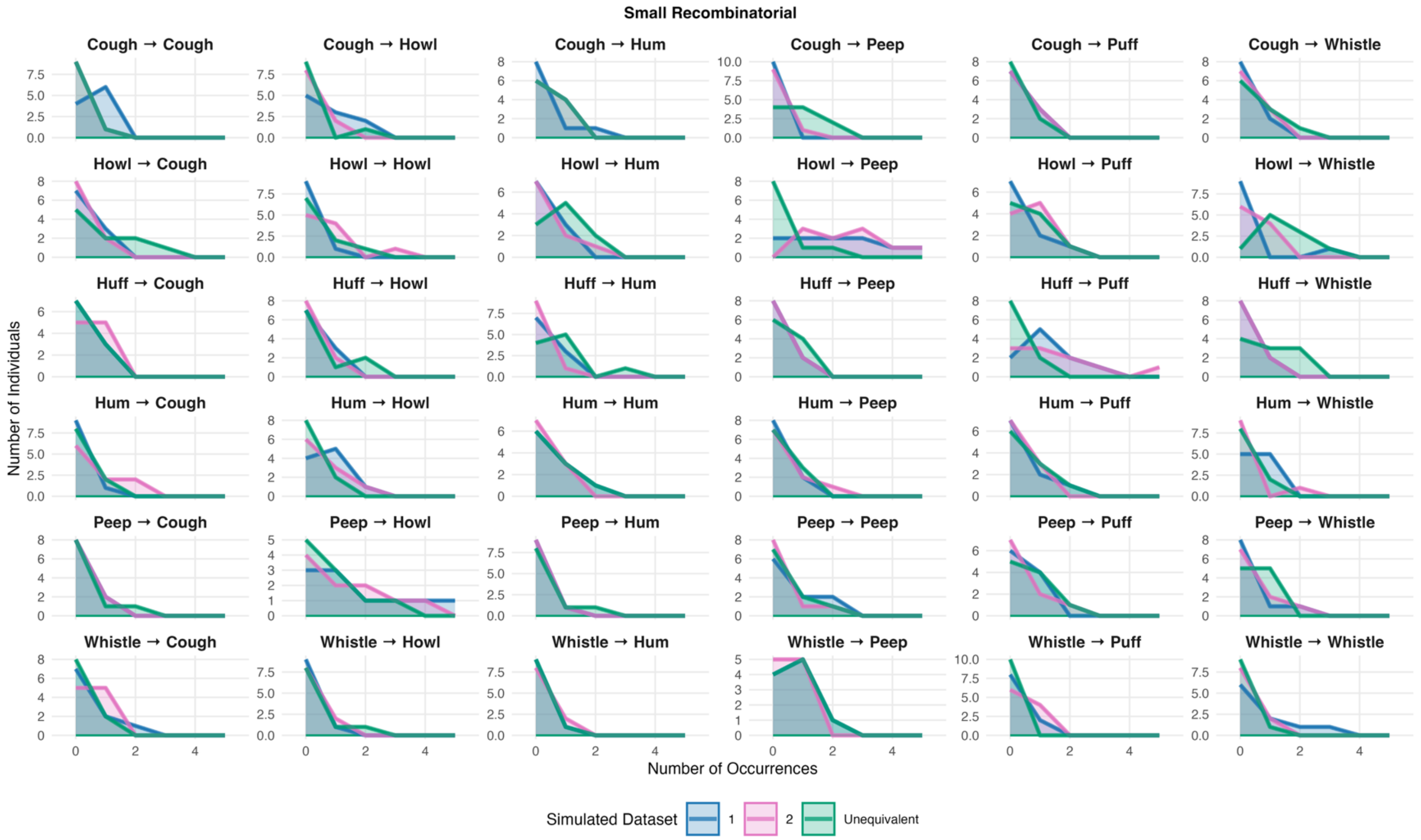

**Fig. S3 Frequency distributions for all bigram types per simulated individual in the *Small Recombinatorial* corpora of Study 2**.

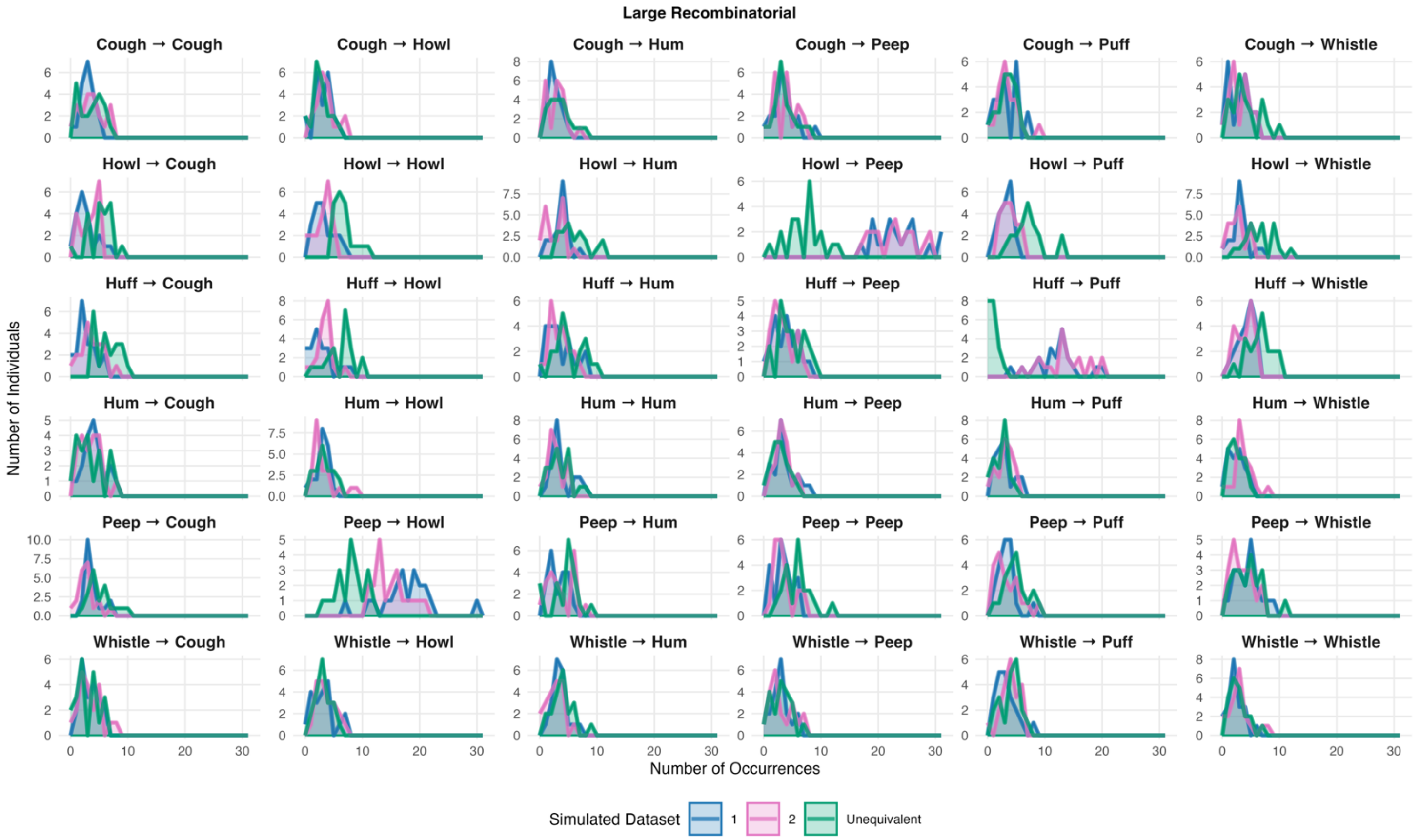

**Fig. S4 Frequency distributions for bigram types per simulated individual in the *Large Recombinatorial* corpora of Study 2**.

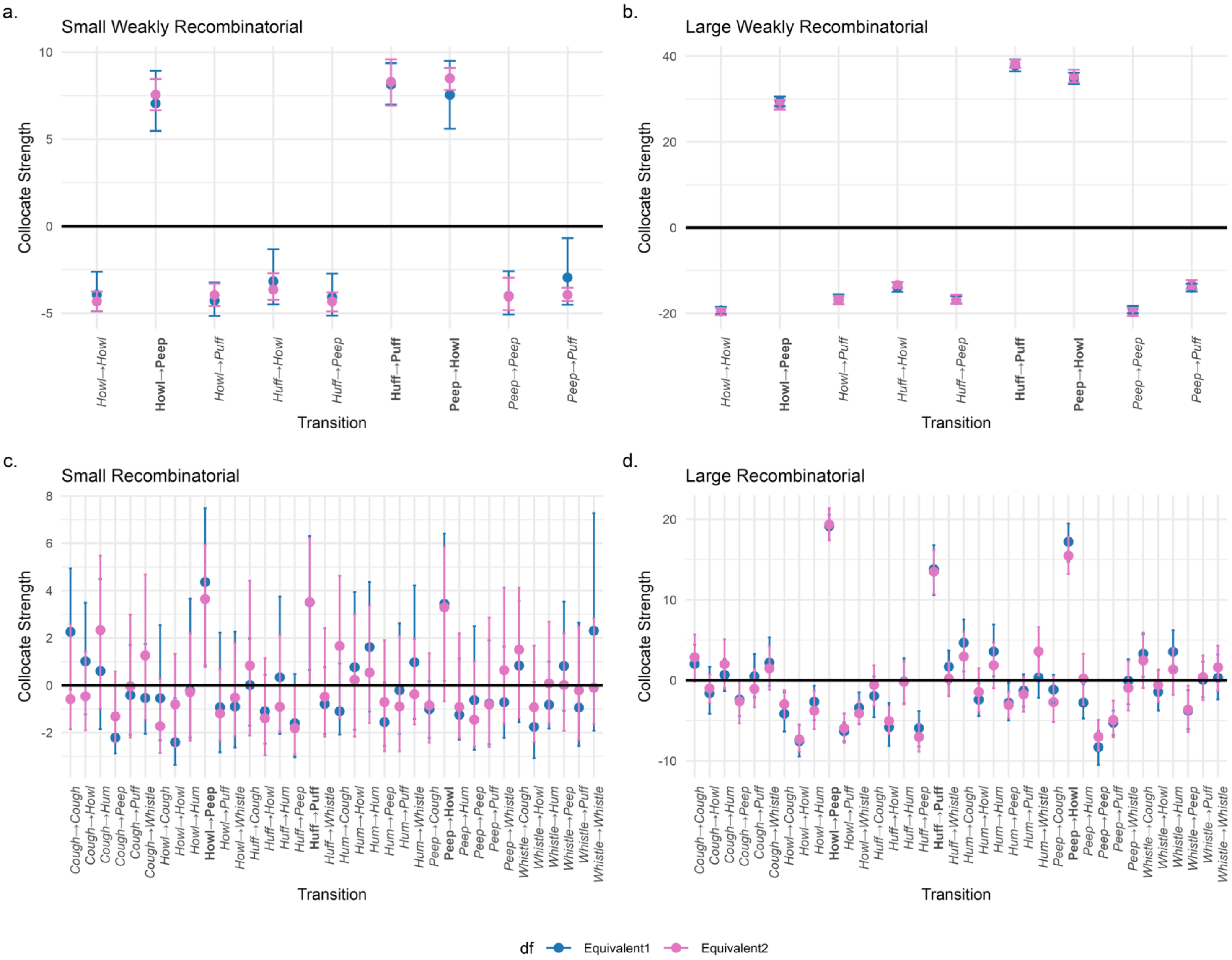

**Fig. S5 MDCA-Pr results for *Equivalent* cohorts 1 and 2 for each data class.**

**Table S5. MDCA-Pr results for comparisons in the *Small Weakly Recombinatorial* data class.**

| Bigram type | Cohort 1 | Cohort 2 | Cohort 1: Lower CI | Cohort 1: Upper CI | Cohort 2: Lower CI | Cohort 2: Upper CI | Overlap |
|---|---|---|---|---|---|---|---|
| Howl→Howl | Equivalent1 | Equivalent2 | -4.891 | -2.610 | -4.87 | -3.745 | TRUE |
| Howl→Peep | Equivalent1 | Equivalent2 | 5.479 | 8.933 | 6.657 | 8.456 | TRUE |
| Howl→Puff | Equivalent1 | Equivalent2 | -5.148 | -3.237 | -4.579 | -3.298 | TRUE |
| Huff→Howl | Equivalent1 | Equivalent2 | -4.489 | -1.327 | -4.228 | -2.696 | TRUE |
| Huff→Peep | Equivalent1 | Equivalent2 | -5.131 | -2.722 | -4.899 | -3.792 | TRUE |
| Huff→Puff | Equivalent1 | Equivalent2 | 6.989 | 9.373 | 6.930 | 9.592 | TRUE |
| Peep→Howl | Equivalent1 | Equivalent2 | 5.599 | 9.498 | 7.827 | 9.099 | TRUE |
| Peep→Peep | Equivalent1 | Equivalent2 | -5.078 | -2.582 | -4.817 | -2.958 | TRUE |
| Peep→Puff | Equivalent1 | Equivalent2 | -4.506 | -0.682 | -4.300 | -3.536 | TRUE |
| Howl→Howl | Equivalent1 | Unequivalent | -4.891 | -2.610 | -4.879 | -2.630 | TRUE |
| Howl→Peep | Equivalent1 | Unequivalent | 5.479 | 8.933 | 3.815 | 6.124 | TRUE |
| Howl→Puff | Equivalent1 | Unequivalent | -5.148 | -3.237 | -2.807 | 1.181 | FALSE |
| Huff→Howl | Equivalent1 | Unequivalent | -4.489 | -1.327 | -2.947 | 1.257 | TRUE |
| Huff→Peep | Equivalent1 | Unequivalent | -5.131 | -2.722 | -3.099 | -0.542 | TRUE |
| Huff→Puff | Equivalent1 | Unequivalent | 6.989 | 9.373 | 1.225 | 6.907 | FALSE |
| Peep→Howl | Equivalent1 | Unequivalent | 5.599 | 9.498 | 2.842 | 6.966 | TRUE |
| Peep→Peep | Equivalent1 | Unequivalent | -5.078 | -2.582 | -5.203 | -2.385 | TRUE |
| Peep→Puff | Equivalent1 | Unequivalent | -4.506 | -0.682 | -3.326 | 0.389 | TRUE |

**Table S6. MDCA-Pr results for comparisons in the *Large Weakly Recombinatorial* data class.**

| Bigram type | Cohort 1 | Cohort 2 | Cohort 1: Lower CI | Cohort 1: Upper CI | Cohort 2: Lower CI | Cohort 2: Upper CI | Overlap |
|---|---|---|---|---|---|---|---|
| Howl→Howl | Equivalent1 | Equivalent2 | -20.195 | -18.487 | -20.422 | -18.726 | TRUE |
| Howl→Peep | Equivalent1 | Equivalent2 | 28.318 | 30.544 | 27.536 | 29.786 | TRUE |
| Howl→Puff | Equivalent1 | Equivalent2 | -17.877 | -15.555 | -17.799 | -15.936 | TRUE |
| Huff→Howl | Equivalent1 | Equivalent2 | -14.975 | -12.721 | -14.060 | -12.724 | TRUE |
| Huff→Peep | Equivalent1 | Equivalent2 | -17.737 | -15.98 | -17.733 | -15.666 | TRUE |
| Huff→Puff | Equivalent1 | Equivalent2 | 36.417 | 39.253 | 37.279 | 39.172 | TRUE |
| Peep→Howl | Equivalent1 | Equivalent2 | 33.482 | 36.141 | 33.835 | 36.832 | TRUE |
| Peep→Peep | Equivalent1 | Equivalent2 | -20.003 | -18.266 | -20.586 | -18.723 | TRUE |
| Peep→Puff | Equivalent1 | Equivalent2 | -14.916 | -13.102 | -14.356 | -12.221 | TRUE |
| Howl→Howl | Equivalent1 | Unequivalent | -20.195 | -18.487 | -14.999 | -13.028 | FALSE |
| How→Peep | Equivalent1 | Unequivalent | 28.318 | 30.544 | 16.958 | 19.658 | FALSE |
| Howl→Puff | Equivalent1 | Unequivalent | -17.877 | -15.555 | -8.204 | -4.414 | FALSE |
| Huff→Howl | Equivalent1 | Unequivalent | -14.975 | -12.721 | -8.105 | -5.724 | FALSE |
| Huff→Peep | Equivalent1 | Unequivalent | -17.737 | -15.98 | -10.023 | -7.221 | FALSE |
| Huff→Puff | Equivalent1 | Unequivalent | 36.417 | 39.253 | 16.930 | 20.921 | FALSE |
| Peep→Howl | Equivalent1 | Unequivalent | 33.482 | 36.141 | 21.157 | 23.947 | FALSE |
| Peep→Peep | Equivalent1 | Unequivalent | -20.003 | -18.266 | -14.727 | -12.731 | FALSE |
| Peep→Puff | Equivalent1 | Unequivalent | -14.916 | -13.102 | -10.820 | -7.346 | FALSE |

**Table S7. MDCA results for comparisons in the *Small Recombinatorial* data class.**

| Bigram type | Cohort 1 | Cohort 2 | Cohort 1: Lower CI | Cohort 1: Upper CI | Cohort 2: Lower CI | Cohort 2: Upper CI | Overlap |
|---|---|---|---|---|---|---|---|
| Cough→Cough | Equivalent1 | Equivalent2 | -0.171 | 4.943 | -1.857 | 2.593 | TRUE |
| Cough→Howl | Equivalent1 | Equivalent2 | -1.230 | 3.484 | -1.901 | 1.401 | TRUE |
| Cough→Hum | Equivalent1 | Equivalent2 | -1.847 | 4.491 | -0.987 | 5.480 | TRUE |
| Cough→Peep | Equivalent1 | Equivalent2 | -2.878 | -1.449 | -2.403 | 0.586 | TRUE |
| Cough→Puff | Equivalent1 | Equivalent2 | -2.098 | 1.700 | -2.214 | 2.984 | TRUE |
| Cough→Whistle | Equivalent1 | Equivalent2 | -2.049 | 1.746 | -1.462 | 4.672 | TRUE |
| Howl→Cough | Equivalent1 | Equivalent2 | -2.324 | 2.550 | -2.853 | 0.282 | TRUE |
| Howl→Howl | Equivalent1 | Equivalent2 | -3.356 | -0.534 | -2.608 | 1.329 | TRUE |
| Howl→Hum | Equivalent1 | Equivalent2 | -2.210 | 3.658 | -2.337 | 2.192 | TRUE |
| Howl→Peep | Equivalent1 | Equivalent2 | 0.839 | 7.480 | 0.763 | 5.938 | TRUE |
| Howl→Puff | Equivalent1 | Equivalent2 | -2.824 | 2.228 | -2.341 | 0.674 | TRUE |
| Howl→Whistle | Equivalent1 | Equivalent2 | -2.634 | 2.257 | -2.136 | 1.792 | TRUE |
| Huff→Cough | Equivalent1 | Equivalent2 | -1.699 | 1.973 | -2.122 | 4.418 | TRUE |
| Huff→Howl | Equivalent1 | Equivalent2 | -2.465 | 0.455 | -2.958 | 1.149 | TRUE |
| Huff→Hum | Equivalent1 | Equivalent2 | -2.055 | 3.752 | -1.970 | 2.121 | TRUE |
| Huff→Peep | Equivalent1 | Equivalent2 | -3.033 | 0.479 | -2.915 | -0.005 | TRUE |
| Huff→Puff | Equivalent1 | Equivalent2 | 0.645 | 6.299 | 0.051 | 6.234 | TRUE |
| Huff→Whistle | Equivalent1 | Equivalent2 | -2.062 | 0.763 | -2.165 | 2.412 | TRUE |
| Hum→Cough | Equivalent1 | Equivalent2 | -2.086 | 0.923 | -1.387 | 4.624 | TRUE |
| Hum→Howl | Equivalent1 | Equivalent2 | -1.884 | 3.942 | -2.162 | 3.000 | TRUE |
| Hum→Hum | Equivalent1 | Equivalent2 | -1.112 | 4.363 | -1.591 | 3.363 | TRUE |
| Hum→Peep | Equivalent1 | Equivalent2 | -2.562 | 0.118 | -2.780 | 1.908 | TRUE |
| Hum→Puff | Equivalent1 | Equivalent2 | -2.054 | 2.614 | -2.786 | 2.097 | TRUE |
| Hum→Whistle | Equivalent1 | Equivalent2 | -1.423 | 4.216 | -1.653 | 1.954 | TRUE |
| Peep→Cough | Equivalent1 | Equivalent2 | -2.225 | 0.171 | -2.428 | 1.357 | TRUE |
| Peep→Howl | Equivalent1 | Equivalent2 | 0.187 | 6.402 | -0.674 | 5.853 | TRUE |
| Peep→Hum | Equivalent1 | Equivalent2 | -2.291 | 1.135 | -2.233 | 2.187 | TRUE |
| Peep→Peep | Equivalent1 | Equivalent2 | -2.725 | 2.491 | -2.598 | 1.024 | TRUE |
| Peep→Puff | Equivalent1 | Equivalent2 | -2.490 | 1.901 | -2.608 | 2.866 | TRUE |
| Peep→Whistle | Equivalent1 | Equivalent2 | -2.225 | 1.637 | -1.609 | 4.124 | TRUE |
| Whistle→Cough | Equivalent1 | Equivalent2 | -1.563 | 3.566 | -1.369 | 4.122 | TRUE |
| Whistle→Howl | Equivalent1 | Equivalent2 | -3.080 | 0.143 | -2.398 | 1.678 | TRUE |
| Whistle→Hum | Equivalent1 | Equivalent2 | -1.823 | 1.006 | -1.612 | 2.688 | TRUE |
| Whistle→Peep | Equivalent1 | Equivalent2 | -1.075 | 3.536 | -1.922 | 2.195 | TRUE |
| Whistle→Puff | Equivalent1 | Equivalent2 | -2.562 | 2.648 | -2.312 | 2.470 | TRUE |
| Whistle→Whistle | Equivalent1 | Equivalent2 | -1.917 | 7.268 | -1.620 | 2.828 | TRUE |
| Cough→Cough | Equivalent1 | Unequivalent | -0.171 | 4.943 | -2.291 | 2.368 | TRUE |
| Cough→Howl | Equivalent1 | Unequivalent | -1.230 | 3.484 | -2.054 | 1.908 | TRUE |

| | | | | | | | |
|---|---|---|---|---|---|---|---|
| Cough→Hum | Equivalent1 | Unequivalent | -1.847 | 4.491 | -2.208 | 2.793 | TRUE |
| Cough→Peep | Equivalent1 | Unequivalent | -2.878 | -1.449 | -1.135 | 6.113 | FALSE |
| Cough→Puff | Equivalent1 | Unequivalent | -2.098 | 1.700 | -1.882 | 1.269 | TRUE |
| Cough→Whistle | Equivalent1 | Unequivalent | -2.049 | 1.746 | -2.068 | 2.946 | TRUE |
| Howl→Cough | Equivalent1 | Unequivalent | -2.324 | 2.550 | -1.703 | 4.417 | TRUE |
| Howl→Howl | Equivalent1 | Unequivalent | -3.356 | -0.534 | -2.683 | 1.688 | TRUE |
| Howl→Hum | Equivalent1 | Unequivalent | -2.210 | 3.658 | -1.408 | 2.746 | TRUE |
| Howl→Peep | Equivalent1 | Unequivalent | 0.839 | 7.480 | -3.376 | 0.538 | FALSE |
| Howl→Puff | Equivalent1 | Unequivalent | -2.824 | 2.228 | -2.204 | 2.684 | TRUE |
| Howl→Whistle | Equivalent1 | Unequivalent | -2.634 | 2.257 | -0.325 | 3.407 | TRUE |
| Huff→Cough | Equivalent1 | Unequivalent | -1.699 | 1.973 | -1.970 | 0.838 | TRUE |
| Huff→Howl | Equivalent1 | Unequivalent | -2.465 | 0.455 | -2.430 | 3.833 | TRUE |
| Huff→Hum | Equivalent1 | Unequivalent | -2.055 | 3.752 | -2.384 | 4.401 | TRUE |
| Huff→Peep | Equivalent1 | Unequivalent | -3.033 | 0.479 | -2.449 | 1.247 | TRUE |
| Huff→Puff | Equivalent1 | Unequivalent | 0.645 | 6.299 | -2.081 | 0.220 | FALSE |
| Huff→Whistle | Equivalent1 | Unequivalent | -2.062 | 0.763 | -1.528 | 2.426 | TRUE |
| Hum→Cough | Equivalent1 | Unequivalent | -2.086 | 0.923 | -2.042 | 3.287 | TRUE |
| Hum→Howl | Equivalent1 | Unequivalent | -1.884 | 3.942 | -2.411 | 3.022 | TRUE |
| Hum→Hum | Equivalent1 | Unequivalent | -1.112 | 4.363 | -1.643 | 4.207 | TRUE |
| Hum→Peep | Equivalent1 | Unequivalent | -2.562 | 0.118 | -1.817 | 1.589 | TRUE |
| Hum→Puff | Equivalent1 | Unequivalent | -2.054 | 2.614 | -1.419 | 4.837 | TRUE |
| Hum→Whistle | Equivalent1 | Unequivalent | -1.423 | 4.216 | -2.469 | 1.008 | TRUE |
| Peep→Cough | Equivalent1 | Unequivalent | -2.225 | 0.171 | -2.156 | 2.345 | TRUE |
| Peep→Howl | Equivalent1 | Unequivalent | 0.187 | 6.402 | -1.933 | 6.008 | TRUE |
| Peep→Hum | Equivalent1 | Unequivalent | -2.291 | 1.135 | -2.529 | 1.789 | TRUE |
| Peep→Peep | Equivalent1 | Unequivalent | -2.725 | 2.491 | -2.482 | 2.089 | TRUE |
| Peep→Puff | Equivalent1 | Unequivalent | -2.490 | 1.901 | -1.521 | 4.023 | TRUE |
| Peep→Whistle | Equivalent1 | Unequivalent | -2.225 | 1.637 | -2.578 | 2.155 | TRUE |
| Whistle→Cough | Equivalent1 | Unequivalent | -1.563 | 3.566 | -1.579 | 3.386 | TRUE |
| Whistle→Howl | Equivalent1 | Unequivalent | -3.080 | 0.143 | -1.725 | 4.608 | TRUE |
| Whistle→Hum | Equivalent1 | Unequivalent | -1.823 | 1.006 | -1.856 | 1.279 | TRUE |
| Whistle→Peep | Equivalent1 | Unequivalent | -1.075 | 3.536 | -0.32 | 6.235 | TRUE |
| Whistle→Puff | Equivalent1 | Unequivalent | -2.562 | 2.648 | -1.791 | -0.908 | TRUE |
| Whistle→Whistle | Equivalent1 | Unequivalent | -1.917 | 7.268 | -2.040 | 0.218 | TRUE |
| Cough→Cough | Equivalent1 | Unequivalent | -0.171 | 4.943 | -2.291 | 2.368 | TRUE |

**Table S8. MDCA-Pr results for comparisons in the *Large Recombinatorial* data class.**

| Bigram type | Cohort 1 | Cohort 2 | Cohort 1: Lower CI | Cohort 1: Upper CI | Cohort 2: Lower CI | Cohort 2: Upper CI | Overlap |
|---|---|---|---|---|---|---|---|
| Cough→Cough | Equivalent1 | Equivalent2 | -0.095 | 4.396 | -0.161 | 5.689 | TRUE |
| Cough→Howl | Equivalent1 | Equivalent2 | -4.145 | 1.664 | -2.746 | 0.720 | TRUE |
| Cough→Hum | Equivalent1 | Equivalent2 | -1.318 | 2.652 | -0.968 | 5.082 | TRUE |
| Cough→Peep | Equivalent1 | Equivalent2 | -4.462 | 0.031 | -5.320 | -0.039 | TRUE |
| Cough→Puff | Equivalent1 | Equivalent2 | -2.088 | 3.268 | -3.338 | 1.392 | TRUE |
| Cough→Whistle | Equivalent1 | Equivalent2 | -0.701 | 5.334 | -1.164 | 4.235 | TRUE |
| Howl→Cough | Equivalent1 | Equivalent2 | -6.347 | -1.473 | -4.826 | -1.223 | TRUE |
| Howl→Howl | Equivalent1 | Equivalent2 | -9.441 | -5.517 | -8.84 | -5.562 | TRUE |
| Howl→Hum | Equivalent1 | Equivalent2 | -4.928 | -0.691 | -6.047 | -1.291 | TRUE |
| Howl→Peep | Equivalent1 | Equivalent2 | 17.415 | 20.599 | 17.393 | 21.367 | TRUE |
| Howl→Puff | Equivalent1 | Equivalent2 | -7.545 | -5.216 | -7.773 | -4.113 | TRUE |
| Howl→Whistle | Equivalent1 | Equivalent2 | -5.382 | -1.477 | -5.474 | -2.519 | TRUE |
| Huff→Cough | Equivalent1 | Equivalent2 | -4.577 | 0.491 | -3.185 | 1.856 | TRUE |
| Huff→Howl | Equivalent1 | Equivalent2 | -8.143 | -2.814 | -6.667 | -3.249 | TRUE |
| Huff→Hum | Equivalent1 | Equivalent2 | -2.868 | 2.756 | -2.982 | 2.393 | TRUE |
| Huff→Peep | Equivalent1 | Equivalent2 | -8.192 | -3.830 | -8.837 | -5.448 | TRUE |
| Huff→Puff | Equivalent1 | Equivalent2 | 10.594 | 16.791 | 10.712 | 16.187 | TRUE |
| Huff→Whistle | Equivalent1 | Equivalent2 | -0.503 | 3.694 | -1.956 | 2.333 | TRUE |
| Hum→Cough | Equivalent1 | Equivalent2 | 1.104 | 7.564 | -0.069 | 6.014 | TRUE |
| Hum→Howl | Equivalent1 | Equivalent2 | -4.464 | -0.413 | -3.915 | 1.507 | TRUE |
| Hum→Hum | Equivalent1 | Equivalent2 | 0.507 | 6.936 | -0.982 | 4.688 | TRUE |
| Hum→Peep | Equivalent1 | Equivalent2 | -4.972 | -0.300 | -4.647 | -1.452 | TRUE |
| Hum→Puff | Equivalent1 | Equivalent2 | -3.295 | 0.770 | -3.907 | 0.438 | TRUE |
| Hum→Whistle | Equivalent1 | Equivalent2 | -2.172 | 2.975 | 0.595 | 6.615 | TRUE |
| Peep→Cough | Equivalent1 | Equivalent2 | -2.797 | 0.669 | -5.175 | 0.519 | TRUE |
| Peep→Howl | Equivalent1 | Equivalent2 | 14.741 | 19.46 | 13.193 | 17.789 | TRUE |
| Peep→Hum | Equivalent1 | Equivalent2 | -4.732 | -0.501 | -3.008 | 3.296 | TRUE |
| Peep→Peep | Equivalent1 | Equivalent2 | -10.478 | -6.266 | -8.719 | -4.910 | TRUE |
| Peep→Puff | Equivalent1 | Equivalent2 | -6.785 | -3.733 | -7.046 | -2.526 | TRUE |
| Peep→Whistle | Equivalent1 | Equivalent2 | -2.962 | 2.608 | -3.727 | 2.404 | TRUE |
| Whistle→Cough | Equivalent1 | Equivalent2 | 0.463 | 5.752 | -0.935 | 5.946 | TRUE |
| Whistle→Howl | Equivalent1 | Equivalent2 | -3.747 | 1.279 | -2.667 | 1.312 | TRUE |
| Whistle→Hum | Equivalent1 | Equivalent2 | 1.204 | 6.239 | -1.800 | 3.990 | TRUE |
| Whistle→Peep | Equivalent1 | Equivalent2 | -6.063 | -1.209 | -6.405 | -0.670 | TRUE |
| Whistle→Puff | Equivalent1 | Equivalent2 | -2.081 | 2.327 | -2.430 | 3.082 | TRUE |
| Whistle→Whistle | Equivalent1 | Equivalent2 | -2.371 | 3.193 | -1.255 | 4.314 | TRUE |
| Cough→Cough | Equivalent1 | Unequivalent | -0.095 | 4.396 | -2.197 | 2.722 | TRUE |
| Cough→Howl | Equivalent1 | Unequivalent | -4.145 | 1.664 | -4.713 | -0.756 | TRUE |

| | | | | | | | |
|---|---|---|---|---|---|---|---|
| Cough→Hum | Equivalent1 | Unequivalent | -1.318 | 2.652 | -2.551 | 2.827 | TRUE |
| Cough→Peep | Equivalent1 | Unequivalent | -4.462 | 0.031 | -2.615 | 3.014 | TRUE |
| Cough→Puff | Equivalent1 | Unequivalent | -2.088 | 3.268 | -1.761 | 4.175 | TRUE |
| Cough→Whistle | Equivalent1 | Unequivalent | -0.701 | 5.334 | -1.235 | 4.084 | TRUE |
| Howl→Cough | Equivalent1 | Unequivalent | -6.347 | -1.473 | -4.155 | -0.290 | TRUE |
| Howl→Howl | Equivalent1 | Unequivalent | -9.441 | -5.517 | -2.489 | 0.964 | FALSE |
| Howl→Hum | Equivalent1 | Unequivalent | -4.928 | -0.691 | -2.952 | 2.081 | TRUE |
| Howl→Peep | Equivalent1 | Unequivalent | 17.415 | 20.599 | -1.940 | 3.703 | FALSE |
| Howl→Puff | Equivalent1 | Unequivalent | -7.545 | -5.216 | 0.680 | 5.331 | FALSE |
| Howl→Whistle | Equivalent1 | Unequivalent | -5.382 | -1.477 | -2.430 | 2.436 | TRUE |
| Huff→Cough | Equivalent1 | Unequivalent | -4.577 | 0.491 | 0.404 | 4.970 | TRUE |
| Huff→Howl | Equivalent1 | Unequivalent | -8.143 | -2.814 | -1.604 | 3.284 | FALSE |
| Huff→Hum | Equivalent1 | Unequivalent | -2.868 | 2.756 | -2.734 | 3.055 | TRUE |
| Huff→Peep | Equivalent1 | Unequivalent | -8.192 | -3.830 | -3.729 | 2.140 | FALSE |
| Huff→Puff | Equivalent1 | Unequivalent | 10.594 | 16.791 | -8.442 | -5.910 | FALSE |
| Huff→Whistle | Equivalent1 | Unequivalent | -0.503 | 3.694 | 0.410 | 5.536 | TRUE |
| Hum→Cough | Equivalent1 | Unequivalent | 1.104 | 7.564 | -2.666 | 4.769 | TRUE |
| Hum→Howl | Equivalent1 | Unequivalent | -4.464 | -0.413 | -2.143 | 1.856 | TRUE |
| Hum→Hum | Equivalent1 | Unequivalent | 0.507 | 6.936 | -1.047 | 4.714 | TRUE |
| Hum→Peep | Equivalent1 | Unequivalent | -4.972 | -0.300 | -2.768 | 1.575 | TRUE |
| Hum→Puff | Equivalent1 | Unequivalent | -3.295 | 0.770 | -3.022 | 2.452 | TRUE |
| Hum→Whistle | Equivalent1 | Unequivalent | -2.172 | 2.975 | -3.236 | 0.322 | TRUE |
| Peep→Cough | Equivalent1 | Unequivalent | -2.797 | 0.669 | -3.374 | 2.151 | TRUE |
| Peep→Howl | Equivalent1 | Unequivalent | 14.741 | 19.46 | 0.652 | 5.946 | FALSE |
| Peep→Hum | Equivalent1 | Unequivalent | -4.732 | -0.501 | -4.330 | 0.131 | TRUE |
| Peep→Peep | Equivalent1 | Unequivalent | -10.478 | -6.266 | -1.994 | 4.280 | FALSE |
| Peep→Puff | Equivalent1 | Unequivalent | -6.785 | -3.733 | -2.224 | 3.254 | FALSE |
| Peep→Whistle | Equivalent1 | Unequivalent | -2.962 | 2.608 | -4.502 | 0.608 | TRUE |
| Whistle→Cough | Equivalent1 | Unequivalent | 0.463 | 5.752 | -3.383 | 1.224 | TRUE |
| Whistle→Howl | Equivalent1 | Unequivalent | -3.747 | 1.279 | -3.431 | 1.333 | TRUE |
| Whistle→Hum | Equivalent1 | Unequivalent | 1.204 | 6.239 | -1.157 | 4.839 | TRUE |
| Whistle→Peep | Equivalent1 | Unequivalent | -6.063 | -1.209 | -2.809 | 1.382 | TRUE |
| Whistle→Puff | Equivalent1 | Unequivalent | -2.081 | 2.327 | -0.135 | 6.051 | TRUE |
| Whistle→Whistle | Equivalent1 | Unequivalent | -2.371 | 3.193 | -3.533 | 1.119 | TRUE |

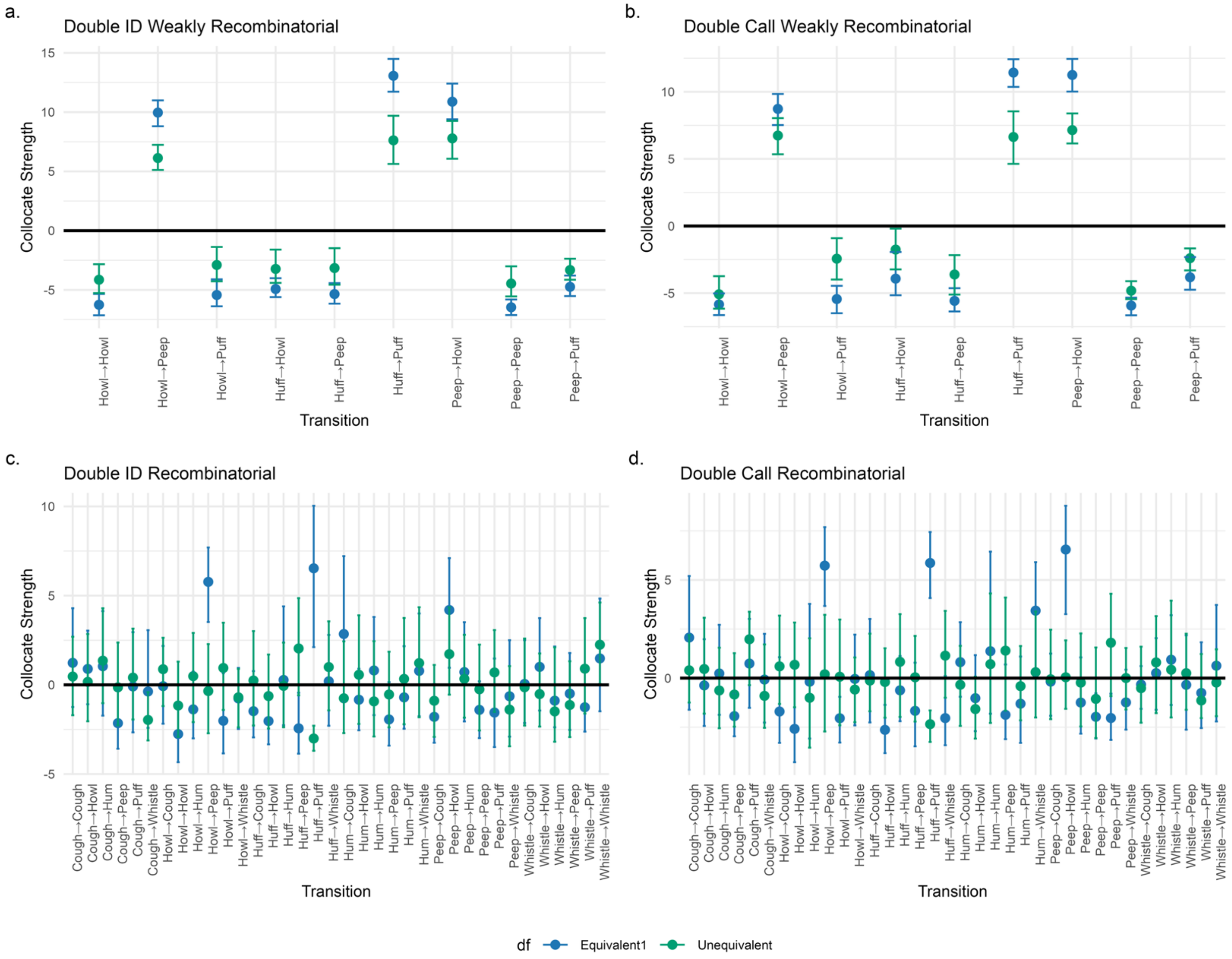

**Fig. S6. MDCA-Pr results for *Equivalent* cohort 1 and the *Unequivalent* cohort after doubling the *Small* sample size.**

**Table S9. Call definitions and number of times sampled in each corpus.**

| Call Type | Description | Sample | |
|---|---|---|---|
| | | With repeats | Without repeats |
| Ekk | ~20-80ms, low intensity ~0.8-1.6kHz, clear harmonic structure (up to 10kHz)[1] ; likely indicates mild anxiety; as tsik-ekk compound call in mobbing or predator alarm | 513 | 234 |
| Food Peep | Short chirping calls, usually uttered in a sequence[2] | 1808 | 105 |
| Food Phee | Short phee at the beginning of/within/at the end of a food peep bout | 89 | 23 |
| Phee | Whistle-like tonal vocalization; 5-9kHz, above 10kHz also possible[1]; typically 1-2s[3] , shorter also possible, but usually > 0.5s; long-distance contact call[4,5]; individually distinct[6]; antiphonal call exchanges[5,7] | 166 | 71 |
| Phee-Peep | Phee-like short vocalization (0.15±0.08sec), low intensity; usually part of a compound call[1] | 93 | 61 |
| Short-Phee | Phee-like call that is between a phee and a phee-like peep in duration | 82 | 59 |
| Trill-Peep | Trill-like short vocalization (0.03-0.2s)[1] (Trills are low-amplitude, frequency-modulated contact calls usually occurring when animals are in close proximity) | 34 | 13 |
| Tsik | Broadband simple call, short duration[1] given in contexts of general anxiety/fear, isolation in unfamiliar environments[8]; seems to have a calming function after stressful situations (decrease in CORT after playback of conspecific tsik calls[9]) | 223 | 221 |

**References for Table S9:**

**1** Agamaite, J. A., Chang, C.-J., Osmanski, M. S., & Wang, X. (2015). A quantitative acoustic analysis of the vocal repertoire of the common marmoset (*Callithrix jacchus*). *The Journal of the Acoustical Society of America*, *138*(5), 2906–2928. https://doi.org/10.1121/1.4934268

**2** Rogers, L. J., Stewart, L., & Kaplan, G. (2018). Food calls in common marmosets, *Callithrix jacchus*, and evidence that one is functionally referential. *Animals*, *8*(7), Article 7. https://doi.org/10.3390/ani8070099

**3** Pistorio, A. L., Vintch, B., & Wang, X. (2006). Acoustic analysis of vocal development in a New World primate, the common marmoset (*Callithrix jacchus*). *The Journal of the Acoustical Society of America*, *120*(3), 1655–1670. https://doi.org/10.1121/1.2225899

**4** Epple, G. (1968). Comparative studies on vocalization in marmoset monkeys (Hapalidae). *Folia Primatologica*, *8*(1), 1–40. https://doi.org/DOI:10.1159/000155129

**5** Miller, C. T., & Wang, X. (2006). Sensory-motor interactions modulate a primate vocal behavior: Antiphonal calling in common marmosets. *Journal of Comparative Physiology A*, *192*(1), 27–38. https://doi.org/10.1007/s00359-005-0043-z

**6** Miller, C. T., Mandel, K., & Wang, X. (2010). The communicative content of the common marmoset phee call during antiphonal calling. *American Journal of Primatology*, *72*(11), 974–980. https://doi.org/10.1002/ajp.20854

**7** Takahashi, D. Y., Narayanan, D. Z., & Ghazanfar, A. A. (2013). Coupled oscillator dynamics of vocal turn-taking in monkeys. *Current Biology*, *23*(21), 2162–2168. https://doi.org/10.1016/j.cub.2013.09.005

**8** Kato, Y., Gokan, H., Oh-Nishi, A., Suhara, T., Watanabe, S., & Minamimoto, T. (2014). Vocalizations associated with anxiety and fear in the common marmoset (*Callithrix jacchus*). *Behavioural Brain Research*, *275*, 43–52. https://doi.org/10.1016/j.bbr.2014.08.047

**9** Cross, N., & Rogers, L. J. (2006). Mobbing vocalizations as a coping response in the common marmoset. *Hormones and Behavior*, *49*(2), 237–245. https://doi.org/10.1016/j.yhbeh.2005.07.007

**Table S10. Marmoset bigram data for each dyad.**

| Dyad | Individual | Sex | Analysis with Repeats | | Analysis without Repeats | |
|---|---|---|---|---|---|---|
| | | | Bigrams | Included? | Bigrams without Repeats | Included? |
| Conan – Mibba | Mibba | F | 77 | Y | 7 | N |
| | Conan | M | 487 | Y | 154 | Y |
| Ginger - Jam | Ginger | F | 245 | Y | 60 | Y |
| | Jam | M | 156 | Y | 29 | Y |
| Jaja - Membo | Jaja | F | 94 | Y | 6 | N |
| | Membo | M | 404 | Y | 15 | N |
| Mojita - Umberto | Mojita | F | 198 | Y | 23 | Y |
| | Umberto | M | 242 | Y | 22 | Y |
| Noccioletta - Ulysses | Noccioletta | F | 469 | Y | 139 | Y |
| | Ulysses | M | 131 | Y | 61 | Y |

**Table S11. MDCA-Pr results on marmoset calls where repeats are included in the data.**

| Sex Category | Call combination | Male | | | Female | | |
|---|---|---|---|---|---|---|---|
| | | Mean | Lower CI | Upper CI | Mean | Lower CI | Upper CI |
| With Repeats | | | | | | | |
| Both Sexes | Ekk→Ekk | 8.419 | 2.885 | 15.521 | 10.223 | 5.956 | 14.071 |
| | Tsik→Ekk | 9.901 | 4.676 | 15.861 | 10.787 | 6.420 | 19.570 |
| | Food Peep→Food Peep | 7.684 | 3.086 | 15.875 | 5.289 | 2.399 | 9.415 |
| | Phee→Phee | 12.490 | 6.151 | 19.163 | 11.191 | 7.026 | 14.575 |
| Male-Biased | Food Phee→Food Phee | 15.707 | 10.975 | 19.061 | - | - | - |
| Female-Biased | Phee→Short Phee | - | - | - | 8.186 | 4.098 | 12.605 |

**Table S12. Post-hoc analysis of attracted call combinations in the underlying data.**

| Sex Category | Call combination | Individuals who produced this transition: Males | Individuals who produced this transition: Females |
|---|---|---|---|
| With Repeats | | | |
| Both Sexes | Ekk→Ekk | Conan: 66<br>Jam: 8<br>Ulysses: 56 | Ginger: 20<br>Noccioletta: 124 |
| | Tsik→Ekk | Conan: 78<br>Jam: 2<br>Ulysses: 48<br>Umberto: 2 | Ginger: 12<br>Jaja: 1<br>Mojita: 3<br>Noccioletta: 71 |
| | Food Peep → Food Peep | Conan: 245<br>Jam: 106<br>Membo: 333<br>Umberto: 182 | Ginger: 147<br>Jaja: 79<br>Mibba: 59<br>Mojita: 165<br>Noccioletta: 172 |
| | Phee → Phee | Conan: 13<br>Jam: 13<br>Membo: 2<br>Ulysses: 14 | Ginger: 18<br>Jaja: 2<br>Mibba: 11<br>Mojita: 6<br>Noccioletta: 9 |
| Male-Biased | Food Phee → Food Phee | Membo: 54 | Jaja: 7 |
| Female-Biased | Phee → Short Phee | Conan: 10<br>Jam: 6<br>Membo: 1 | Ginger: 15<br>Jaja: 1<br>Mibba: 2<br>Mojita: 6<br>Noccioletta: 3 |
| Without Repeats | | | |
| Both | Tsik → Ekk | Conan: 78<br>Jam: 2<br>Ulysses: 48<br>Umberto: 2 | Ginger: 12<br>Mojita: 3<br>Noccioletta: 71 |
| | Food Phee → Phee | Jam: 3 | Ginger: 2 |
| Female-Biased | Ekk → Tsik | Conan: 13<br>Ulysses: 11 | Ginger: 2<br>Noccioletta: 11 |
| | Phee → Short Phee | Conan: 10<br>Jam: 6 | Ginger: 15<br>Mojita: 6<br>Noccioletta: 3 |
| | Food Peep → Food Phee | - | Ginger: 1 |

### Marmoset results with repeating calls

#### Method

The corpus size for each call type can when repeated calls were included can be found in Table S9. Following the same method outlined in Section 2.3.1, we sampled with replacement an equal number of individuals for either sex for each bootstrap (N= 5 per sex), and subsampled 77 call bigrams per individual (without replacement) to ensure that more voluble individuals were not overrepresented within the sample. When repeats were included, each bootstrap of the dataset had a sample size just over double that of the *Small* sample in Study 2 (385 call bigrams per sex).

#### Results

We identified six significantly attracted call combinations produced by marmosets within feeding contexts (see Table S11). Four attracted call combinations were shared by both sexes: repetitions of long-distance Phee calls; repetitions of short-distance Food Peeps; and two call combinations produced by marmosets when in negatively-aroused contexts, including repetitions of Ekk calls, and combinations of Tsik→Ekk calls (Agamaite et al. 2015; Bosshard et al. 2024). For all attracted call combinations shared between the sexes, we identified no sex differences in collocation strength. Additionally, post hoc analysis of these call combinations revealed that each call combination was produced by a similar number of individuals of either sex, and by most individuals sampled (see Table S12).

One male-biased call combination was identified, which was the repetition of long-distance Food Phees. Post hoc analysis revealed these repetitive Food-Phees identified for male marmosets were produced by one male (Membo) who produced 54 repetitions of Food Phees. One female marmoset produced seven Food-Phee repetitions, indicating that this call combination was present in the female data although much less frequently. Given that the many Food-Phee repetitions in the male data were produced by a single individual, we cannot rule out that Food-Phee repetitions were an individual-biased call combinations, rather than being a behaviour typical to all males.

One female-biased call combination was identified, where long-distance Phee calls where combined with a shorter version of the call: a Short Phee. This reflected the result we discuss within the main text of our manuscript, where we compare transitions between unique call types only (see section 2.3.3).

**Comparison with results where repeats are removed**

We therefore found some differences in our results when repeats are included in the analysis of call sequence data by MDCA-Pr. The choice of whether to include repeated calls within the analysis likely depends on the communication system under study. However, we recommend that the methodological consequences of including repeats be considered prior to applying MCDA-Pr. For species with high frequencies of repetitions, these repetitions may mask rarer combinations of different call types which are important for communication and behavioural coordination.
